# Self-organized emergence of modularity, hierarchy, and mirror reversals from competitive synaptic growth in a developmental model of the visual pathway

**DOI:** 10.1101/2024.01.07.574543

**Authors:** Sarthak Chandra, Mikail Khona, Talia Konkle, Ila R. Fiete

**Affiliations:** Department of Brain and Cognitive Sciences & McGovern Institute, MIT; Department of Physics, MIT; Department of Psychology, Harvard University

## Abstract

A hallmark of the primate visual system is its *architectural organization* consisting of multiple distinct (modular) areas that connect hierarchically. These areas exhibit specific *spatial organization* on the cortical sheet, with primary visual cortex at the center and subsequent regions in the hierarchy encircling the earlier one, and detailed *topological organization*, with retinotopy in each area but striking mirror reversals across area boundaries. The developmental rules that drive the simultaneous formation of these architectural, spatial, and topographic aspects of organization are unknown. Here we demonstrate that a simple synaptic growth rule driven by spontaneous activity and heterosynaptic competition generates a detailed connectome of the visual pathway, with emergence of all three types of organization. We identify a theoretical principle — local greedy wiring minimization via spontaneous drive (GWM-S) — implemented by the mechanism, and use this insight to propose biologically distinct growth rules that predict similar endpoints but testably distinguishable developmental trajectories. The same rules predict how input geometry and cortical geometry together drive emergence of hierarchical, convolution-like, spatially and topographically organized sensory processing pathways for different modalities and species, providing a possible explanation for the observed pluripotency of cortical structure formation. We find that the few parameters governing structure emergence in the growth rule constitute simple knobs for rich control, that could (potentially genetically) encode a projection neuron-like connectivity patterns and interneuron-like ones. In all, the presented rules provide a parsimonious mechanistic model for the organization of sensory cortical hierarchies even without detailed genetic cues for features like map reversal, and provide numerous predictions for experiment during normal and perturbed development.

---

The visual cortex of primates consists of multiple discrete visual areas. Each area provides full coverage of the visual field, and the areas process information sequentially and hierarchically. The scale and semantic content of features increases along the hierarchy. Areas in the early visual cortex such as V1, V2, V3, V4, and some areas in the lateral occipital cortex preserve the global topography of visual inputs to the retina [1, 2, 3, 4, 5, 6, 7]. This mapping of nearby locations on the retina to nearby regions of cortex, or retinotopy, is similar to the topographical mapping of inputs seen in other sensory cortices [8, 9, 10]. These hierarchically connected early visual areas must somehow be fit into one two-dimensional cortical sheet while also preserving topographic order. In primates, this is accomplished by spatially organizing areas in a concentric way such that V2 surrounds V1, and so on [11].

In mammals from mice to humans, all three kinds of order — hierarchical, spatial, and topographic — are present before eye-opening or in utero, prior to the receipt of detailed visual stimuli and even before photoreceptors are present [12, 13, 14, 15, 16]. These and other observations suggest that genetic cues and internally generated activity, rather than detailed external visual inputs, drive synaptic wiring. These factors lead to the formation of discrete visual areas and their hierarchical, spatial and topographic organization during development [13].

Developmental biology is spanned by two polar hypotheses. One states that genes directly specify the detailed endpoint architecture of organisms and brains (that brain regions are discrete, how many there are, and their hierarchical connectivity), spatial layout (which part of the cortical sheet the regions occupy and the locations and shapes of their boundaries), and topography (the nature of retinotopic order). This aligns with the chemoaffinity hypothesis of Roger Sperry [17], the positional hypothesis of Lewis Wolpert [18] and extensive experimental work showing a role for genetic hard-wiring or specification [19, 20, 21, 22, 23, 24]. The opposite hypothesis is that the brain is a highly plastic system, a blank slate whose organization is created and shaped by external inputs. This aligns with Karl Lashley’s doctrine of mass action [25], Turing’s pattern formation hypothesis [26], and experimental work showing the flexible process of digit formation [27] and the potential of any region of cortex to develop the properties of another with the right inputs [28].

In this work, we espouse a third hypothesis aligned with the body of work in modern developmental biology showing a middle-path between these poles: in which genes specify a very small number of parameters [29, 30, 31, 32, 33] of a developmental growth process or program, and rich structure then gradually emerges via the unfolding over time of this dynamical process [34, 35, 36]. In other words, structure is created from a dance involving small amounts of genetic information interacting with a largely emergent and self-organizing dynamical process, such that neither genetic information nor external inputs perform endpoint specification. The process involves internally generated spontaneous activity and states that feed back into the unfolding dynamics and serve as context for the next steps of emergence [37]. This view is supported by a body of experiments on cell differentiation, embryogenesis, and in the specific case of the visual system the development of the topographic retinotectal projection [13, 38]. The hypothesis of a bare-bones genetic scaffolding that sets a few parameters of a developmental program, which then unfolds in an activity-dependent way, also opens the door to explaining the fascinating pluripotency of cortical development: that when driven by visual inputs, auditory cortex develops signatures of visual processing [28].

We explore how much brain structure can be induced by the unfolding of simple synaptic growth rules from stimulus-independent spontaneous activity. We consider that competition between neurons and synapses to innervate targets is a cornerstone mechanism for self-organization within and across areas in cortex [39, 40, 41]. Similar competitive dynamics have been shown to organize the one-to-one precise and efficient neural control of muscle fibers at the neuromuscular junction [42, 43, 44] and the crystalline organization of the cerebellum such that multiple climbing fibers that initially innervate cerebellar Purkinje cells are winnowed to a single winner [45]. Competitive models are also consistent with the emergence of sequential premotor neural activity in songbird HVC [46, 47] and with rodent cortex [48]. Finally, the mammalian visual system has provided some of the earliest and continuing evidence of the role of competitive innervation processes[49, 50, 13, 51]. Though our focus in on mammalian circuit development, there is mounting evidence that even in the fly, an ideal of the chemoaffinity hypothesis, there is a critical role for the self-organization of structure driven by spontaneous activity [32, 38, 30, 52, 53].

We find that a simple learning rule, based on presynaptic activity-dependent synaptic growth and heterosynaptic competition[46] can organize an unstructured cortical sheet into a set of discrete regions that are hierarchically connected, with concentric spatial organization on the cortical sheet. Moreover, the topographic order of regions within the hierarchy combines the topography of the inputs with the geometry of the cortical sheet. In the visual input case the result is retinotopic order with mirror reversals observed in topography at region boundaries [9, 8, 1]. Further, the resulting connectivity between adjacent regions in the hierarchy naturally yields a multiscale receptive field structure, with emergent signatures like the eccentricity-dependence of visual receptive fields and local interneuron connectivity coexisting with projection neurons that form the hierarchy. The same rules, applied to cortical sheets with different geometry or different input modality, can explain the organization of visual cortex in different species and the hierarchical organization of various sensory modalities. Finally, we show that this mechanistic process arrives at an endpoint of a local, greedy wiring connectome — offering an alternative to prevalent global wiring minimization theories of cortical circuits [54].

In sum, the model bridges vastly different levels, from a small key set of putatively genetically specified parameters and the subcellular dynamics of synaptic growth and competition, to the mesoscopic organization of maps within cortical regions, to the macroscopic organization of hierarchies between areas. As a result, it provides multiple testable predictions for molecular neuroscience including transcriptomics; developmental neuroscience; and connectomics.

## Results

### Initial architecture, retinal activity, and competitive activity-dependent synaptic growth

The ingredients of our model are: 1) Spontaneous activity in the retinal inputs, putatively driven by retinal waves. 2) Topographic putative thalamic layout with all-to-all (non-topographic) projections into an undifferentiated sheet of cortical neurons [55] with (non-topographic) all-to-all cortico-cortical connectivity. 3) A synaptic growth rate determined by presynaptic activity and soma-to-synapse distance, with heterosynaptic competition between synapses at each presynaptic and postsynaptic neuron [46].

#### Initial model architecture and inputs

We focus on the development of one hemisphere of the visual cortex, and thus restrict the retinal inputs to one visual hemifield, Fig. 1a. Retinal drive, which is retinotopically organized in its projection to an idealized LGN, is scaled in area by the log-conformal transformation [56, 57], Fig. 1b. To start with minimal assumptions about pre-existing connectivity structure in cortex, the connectivity from this idealized LGN to the undifferentiated and unparcellated cortical sheet [55] and within the cortical sheet is set as all-to-all, Fig. 1b. Thus, there are no initial area divisions, topography, or hierarchy in cortex. (We will investigate other versions of unstructured initial connectivity later.)

**Figure 1:**
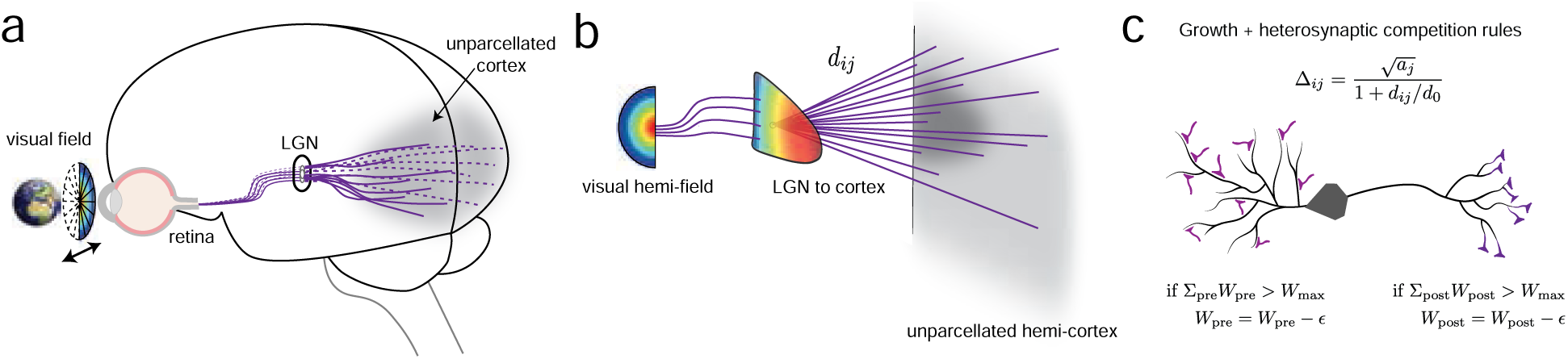
Initial architecture, and competitive synaptic growth rules: (a) A schematic of the visual pathway in primates, wherein visual information is acquired by the retina, and is relayed through the Lateral Geniculate Nucleus(LGN) to reach the visual cortex. We construct our initial architecture based on this structure, beginning with an initially unparcellated cortex. (b) A schematic of our setup, where the visual hemifield of visual input is mapped onto an area representing the LGN. This area is then assumed to lie above and project to a sheet of latent cortical neurons. (c) Top: Synapses grow at a rate that depends on presynaptic activity (*a_j_*) and inversely with the soma-dendritic distance (*d_i j_*). Bottom: The heterosynaptic competition rule: If the sum of incoming (resp. outgoing) weights exceeds a bound (*W_max_*), all synapses undergo depression, a rule that proportionally penalizes weak synapses more than strong ones. Each individual synapse also has a weight bound *w_max_* < *W_max_* (not depicted).

#### Spontaneous activity

Over development, the retina exhibits spontaneous activity in the form of retinal waves [58, 59, 60, 61] that can drive activity in LGN and V1[58, 62]. These inputs have been linked to the formation of the initial visual map before eye-opening [13, 63, 59, 64], and disruptions lead to some visual deficits [65].

We assume that the propagation of retinal activity waves is relatively fast (~10’s of seconds [58]) while synaptic growth and decay is slow (minutes to days [66]) and depends on activity only through the magnitude of presynaptic activity. Therefore, for the current model the spatio-temporal dynamics of the retinal wave patterns are unimportant and on the timescale of plasticity we can replace retinal activity by its average value, which will look like wide-field excitation.

### Synaptic growth and competition rules

We consider a model of synaptic growth driven by presynaptic activity, consistent with experiments [67, 68, 43, 69], Fig.1c. Critically, we assume that the growth rate of a synapse is inversely modified by a distance variable: for cortico-cortical synapses, this distance takes the form of actual soma-synapse distance along the neural arbors, putatively because of costs associated with resource trafficking; for LGN-cortical synapses, this might be an effective distance set by a tract path (e.g. axons must travel along a direct LGN-occipital cortex tract and then to innervate other parts of occipital cortex must grow further). The rate of growth of synapse *i j*, from neuron *j* to neuron *i*, is proportional to Δ*_i j_* :

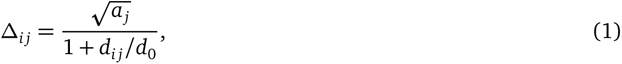

where *d_i j_*/*d*_0_ is a relative soma-to-synapse distance normalized by a length-scale parameter *d*_0_, and *a_j_*is the presynaptic firing rate.

Synapses compete with each other pre- and post-synaptically to drive the postsynaptic neuron [43, 69, 46]: whenever the summed weights into or out of a neuron exceed a threshold (*W*_max_), all the weights into or out of that neuron, respectively, are decremented by a small absolute amount ε, Fig. 1c. This rule is highly competitive, in that small weights are penalized by a larger fraction than large ones, resulting in sparse connectivity with a few strong weights dominating. Each synapse is individually capped at a strength *w*_max_ (< *W*_max_), and weights are kept non-negative, thus distributing connections across roughly *k* = *W*_max_/*w*_max_ synapses per neuron. These bounds might be different for incoming versus outgoing connections and across neurons, but for simplicity we set them to be identical unless otherwise specified. Note that the “weights” grown in the model represent a developmentally developed skeleton, that can be later refined and trained based on visual inputs.

Activity propagates across synapses of strength greater than a threshold of ε*_w_ w*_max_. Neural activity propagates linearly across synapses satisfying this threshold, with the magnitude of activity attenuated by a factor γ for each synapse crossed (see *methods* for details).

### Emergence of a hierarchy of discrete areas with concentric spatial arrangement

Simulating the competitive growth dynamics with spontaneous retinal activity, starting from an unstructured input projection (one-to-all projections) and a maximally unstructured cortical sheet (all-to-all connectivity), Fig. 2a, results in a network with a cellular-resolution synaptic connectome that exhibits remarkable self-organized emergence of order. Initially weak weights strengthen based on distance, so that the shortest-distance connections are strengthened while the heterosynaptic competition penalizes the weaker (and thus longer-distance) ones. Cortical neurons receiving the shortest-distance inputs from LGN gain the strongest weights earliest. As a result, the inputs to these neurons mature fastest (by hitting their saturated values of 0, *w*_max_), and their inputs from elsewhere in cortex weaken. These neurons are the most active, and most effectively able to saturate the weights of their postsynaptic partners, for their closest partners. The strongly competitive process sends most connections to zero, an effective pruning strategy.

**Figure 2:**
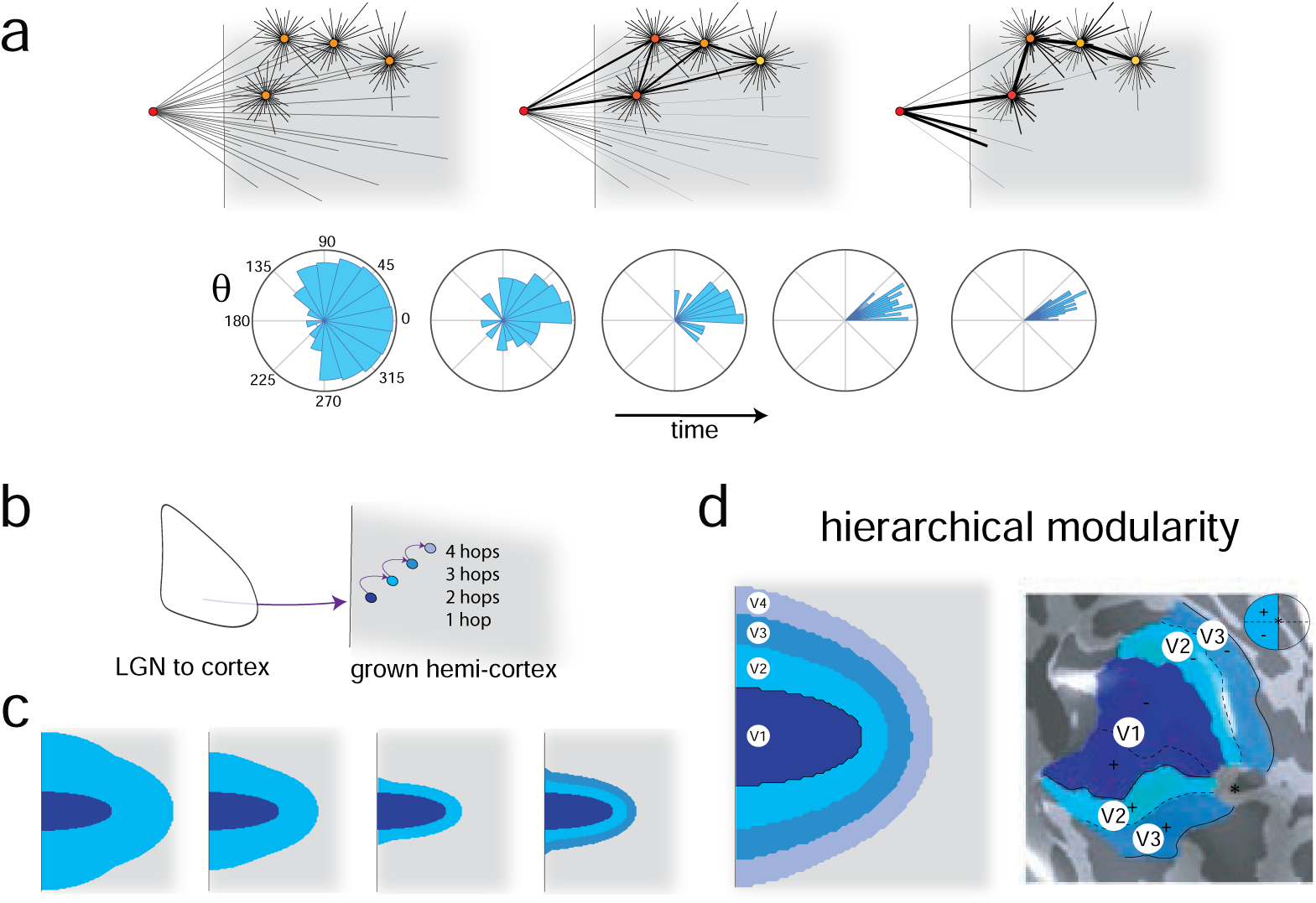
The emergence of a hierarchical connectome from biologically plausible competitive growth rules: (a) Beginning with all-to-all potential recurrent connectivity within cortex and from the putative LGN to cortex, the heterosynaptic competition rule (Fig. 1c) results in the pruning of synapses (top). The angular distribution of postsynaptic target neurons for each neuron in the network grows more directional (bottom). (b) In the formed network, neuron hierarchy can be assigned on the basis of the smallest number of hops with which they are connected to LGN.(c) The heterosynaptic competition growth, based on presynpatic activity, proceeds with each area forming a large number of initial connections (as seen as the wide light-blue shaded area) which are then pruned, eventually leading to the spontaneous emergence of another succeeding area (as determined by the connectivity, (b)).(d) The fully grown and stabilized network forms a set of discrete areas that abut and encircle each other (left), which matches the spatial layout of early visual cortical areas in humans [1](right)

The distribution of outgoing weights of neurons in the cortical sheet becomes increasingly focused and directional, Fig. 2a (bottom). We characterize neurons based on the smallest number of hops with which they are connected to the LGN, Fig. 2b. Initially, all neurons are a single hop from LGN, but as the simulation proceeds, Fig. 2c, most neurons lose single-hop LGN connectivity because the nearest cortical neurons to LGN monopolize the LGN outputs. The outgoing weights from the strongly driven single-hop cortical neurons drive their closest neighbors, to render them two-hop neurons, and so on. As the synaptic dynamics proceed, the neurons stratify into a clear hierarchy of independent discrete regions: neurons that are a single hop away from LGN, two hops away from LGN and so on, Fig. 2d (left). We simulated these dynamics until four such discrete regions emerged (Fig. 2d (left); continued simulation resulted in the formation of additional areas until the available cortical sheet was exhausted, SI Fig. S1).

The formed regions are spatially organized on the cortical sheet in a concentric arrangement: the 4-hop region surrounds the 3-hop region, which encircles the 2-hop region; the 1-hop region is at the center. We putatively call these regions V1, V2, V3, and V4, respectively, Fig. 2d (left). This arrangement of V1, V2 and V3 in our model closely resembles human cortex, in particular in the posterior-medial part of the occipital lobe[1, 2], Fig. 2d (right).

The size of each area is controlled by the ratio of number of incoming to outgoing connections, which is approximately given by the ratio of the total incoming synaptic threshold to the total outgoing synaptic threshold 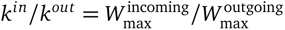 SI Fig. S2.

### Emergence of retinotopy and polar angle alternation

To determine whether there is topographic order in each area of the formed cortical hierarchy, we use standard receptive field mapping procedures, Fig. 3a, [1, 70]. Localized pulses of input in the retinal space drive activity in downstream neurons. Each neuron in the network is then colored based on its preferred retinal location, using different colorings to visualize the mapping of radial eccentricity and polar angle, Fig 3b-c.

**Figure 3:**
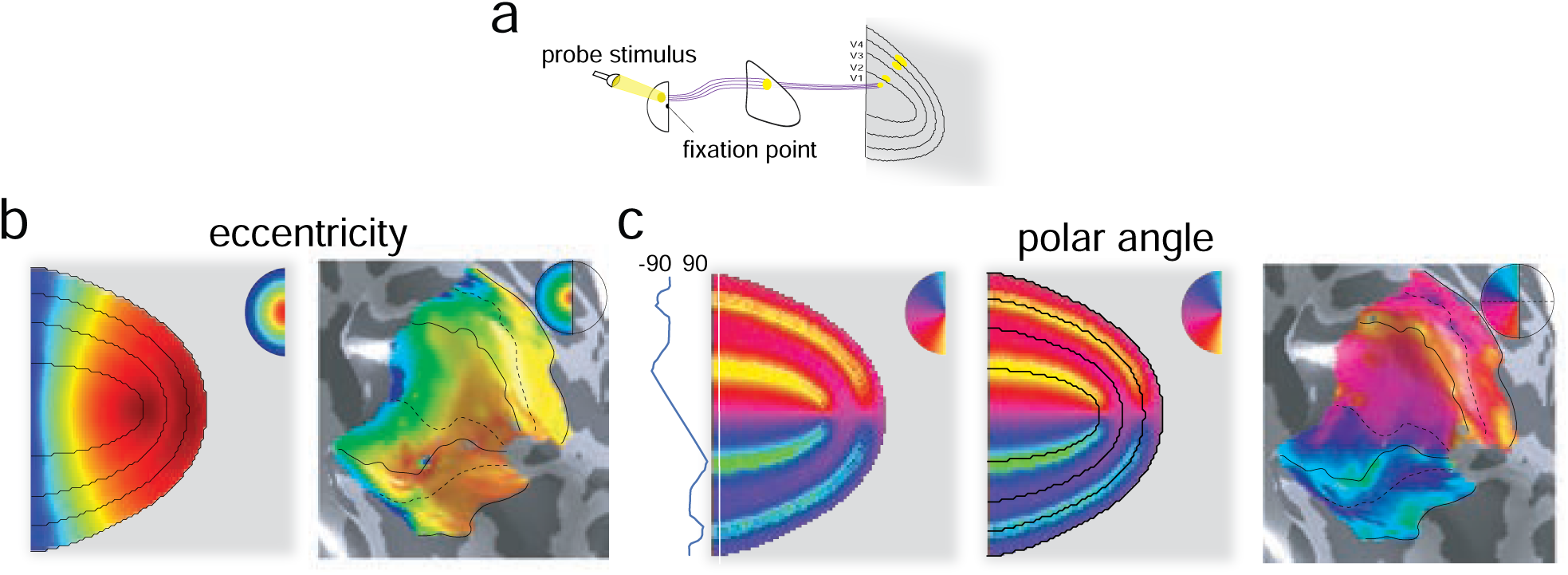
The emergence of mirrored map topography from biologically plausible competitive growth rules: (a) Probing network organization to characterize which neurons respond to point stimuli in the visual field. (b) Neurons on the cortical sheet, colored by their response as a function of radial distance from the fixation point, generate an *eccentricity map* (left), with the corresponding map from human experiments [1] (right). (c) *Polar angle map* (angles from π/2 on the lower vertical meridian up to π/2 on the upper vertical meridian with 0 at the horizontal meridian (left)). Note the alternations in polar angle, in contrast to the smooth eccentricity map. Connectomically defined emergent area boundaries based on the number of hops from LGN (cf. Fig. 2b,d) appear to exactly coincide with the emergent polar angle reversal locations. These reversals occur at the representations of the horizontal and vertical meridians. Polar angle alternations measured in humans [1] (right).

We find that retinal/LGN eccentricity is preserved in the mapping to cortex, within each area and between areas: there is a globally smooth and monotonic mapping from retinal eccentricity to eccentricity representation in the cortex, Fig. 3b (left). More-foveal regions map to one part of the cortical sheet, with an eccentricity gradient that is similar across areas and runs along (parallel to) the area boundaries. This topography in eccentricity, with the log-conformal transformation in LGN, closely matches recordings from human cortex, Fig. 3b (right) [1, 57]. The representation of polar angle is different: Each brain region exhibits local topography for polar angle, but the 90–270*^◦^* polar angle representation is split in two halves, represented by the upper and lower halves of each of the horseshoe-shaped visual area. In contrast to the eccentricity map, polar angle gradients run orthogonal to area boundaries and there is not a globally smooth topographic mapping of polar angle across the hierarchy of areas, Fig. 3c. Polar angle representations exhibit striking mirror reversals at the region boundaries.

This emergent result also matches the findings from human visual cortex, Fig. 3c [1]. Indeed, mirror reversals in polar angle are used to determine area boundaries in many experimental settings [1]. In the model, the independent measure of area boundaries based on hops from LGN aligns perfectly with mirror reversal locations. We believe that this is the first synapse-level mechanistic model of architectural, spatial, and topographic structure emergence in primate visual systems, including the emergence of discrete areas with hierarchical architecture, spatial organization with concentric areas on a cortical sheet, and topographic organization of eccentricity with polar angle reversals coincident with area boundaries. It provides a possible mechanism for the mysterious origin of polar angle alternations, for which there are no known underlying genetic gradient signals [71].

### Distillation of a principle for structure emergence: local greedy wiring minimization

The potent structure-inducing effects of the growth with heterosynaptic competition rule (Eq. 1) suggest the possibility of a general principle at work. Conceptually, spontaneous retinal activity drives the growth of output connections from LGN to cortex. Distance-dependent growth (Eq. 1) promotes more-rapid strengthening of short-distance connections to cortex and within cortex. Simultaneously, the presynaptic activity-dependent growth term means that LGN inputs to cortical cells outcompete cortical inputs to cortical cells. Together, these two effects result in the shortest-distance LGN-to-cortex connections strengthening relative to and at the cost of all other LGN-cortical and cortico-cortical connections. In this way, the very strongly LGN-driven set has become V1, an area strongly innervated by LGN, Fig. 4a (left).

**Figure 4:**
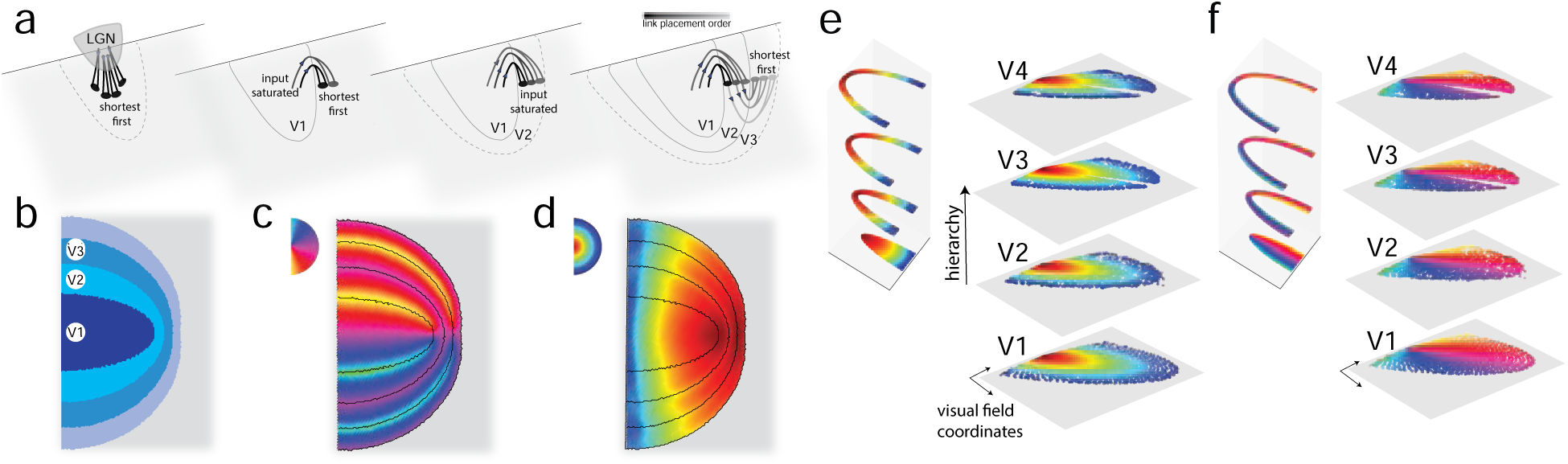
Theoretical principle: Local greedy wiring minimization rule for the emergence of multiple mirrored hierarchical visual areas: (a) A schematic of the greedy wiring minimization principle (left to right): synapses with presynaptic neurons in the previous completely formed area are placed in order of increasing wiring length until all presynaptic neurons saturate in out-degree. The postsynaptic neurons form the next area, and the process continues for this newly formed area. Color denotes the time at which the synapse stabilized. (b-d) The visual hierarchy grown according to the process highlighted above shows the formation of discrete areas with concentric spatial organization of these areas, and mapping (b), polar angle reversals (c) and topographic eccentricity representation (d). (e) Each neuron in the cortical sheet is placed along the z-axis according to its position in the formed hierarchy (left), then positioned on the x-y plane according to the center of its receptive field in visual field coordinates (right). Neurons are colored by eccentricity in (e) and by polar angle in (f).

These V1 neurons have an advantage over all other cortical neurons as potential presynaptic partners to other cortical neurons, because of their higher (LGN-driven) activity. Within this V1 set, neurons closest to the edge of the set have an advantage, because their connections to the surrounding cortical pool are the shortest and strengthen first. Neurons in the interior of V1 are left to strengthen their synapses to available cortical neurons further away. In this way, neurons at the V1 edge outcompete others to innervate the neurons nearest the edge, and neurons at the V1 core connect the furthest away, Fig. 4a (second panel). This is the basis for topographic mirror reversals. As the outputs of V1 saturate and inputs to the edge-proximal target population become strong, this population gains an edge in strengthening its outputs to the remaining cortical neurons, and the next area and mirror-reversal take shape, Fig. 4a (remaining panels).

These insights allow us to hypothesize that the dynamics induced by the learning rule can be distilled into a simple and mathematically principled process that we call *local greedy wiring minimization via spontaneous drive (GWM-S for short)*: starting from a progenitor set of neurons with some topographic ordering of features, in our case the LGN, and a pool of potential target neurons (in our case the undifferentiated cortex), let the progenitor neuron with the closest potential target create a connection at maximal strength to that target. Place the next-nearest connection next (it might belong to the same or another progenitor neuron or end at the same or another target neuron), and so on. Each neuron can place and receive a maximum of *m* connections. Once the progenitor set is saturated, the most-strongly innervated set becomes the progenitor set, and the connection placement process repeats, until all cells are saturated or until cells in the target pool have run out (pseudocode and further detail in the *Methods*). The process is *local* because growth at any point is focused on a subset of cells (growth does not happen globally); it is *greedy* because it proceeds through placement of the shortest available connections at the current moment and from the current set, rather than performing absolute wiring length minimization on the global circuit (as considered in earlier works [72, 54, 73, 74]).

Running the GWM-S process on our initial network recapitulates the architecture, spatial organization, and topography that result from the neural growth and heterosynaptic competition rules, Fig. 4b-d. Thus, neural growth with heterosynaptic competition is a biologically plausible mechanistic process that implements the principle of GWM-S.

### Three conceptually similar but biologically distinct rules reproduce visual hierarchy emergence

The distillation of structure emergence dynamics into the GWM-S process suggests that other growth rules with the same central elements but distinct biological mechanisms might achieve a similar outcome. The central elements are:

1. A seed subset of topographically organized neurons with spontaneous activity.
2. Presynaptic activity-based and distance-dependent synapse formation, strengthening and pruning.
3. In-out-degree saturation in individual neurons.
4. A separation of time-scales between retinal/synaptic activity propagation (fast) and synaptic plasticity (slow).

Indeed, at least two other biologically distinct and plausible growth rules result in quantitatively and qualitatively similar endpoint connectomes, each with hierarchical, spatially concentric, and topographically organized and mirror-reversing retinotopic maps (Fig. 5).

**Figure 5:**
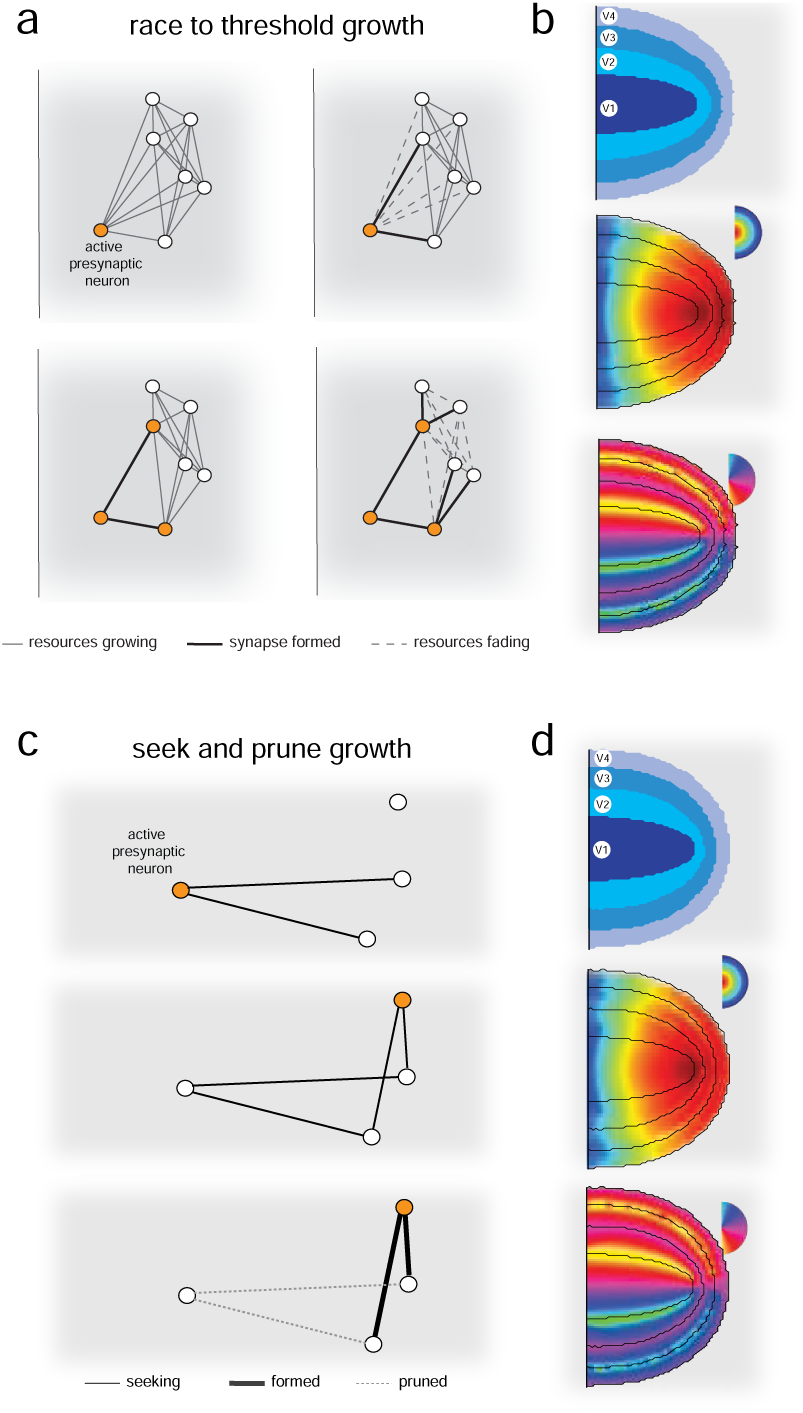
Distinct biologically-plausible learning rules based on growth and pruning reproduce hierarchical, spatial, and topographic organization. (a) The *Race-to-threshold* growth involves the ferrying of resources from active neurons to all their potential synapses. Synapses become active upon crossing a minimum resource thresholds. This resource threshold is higher for further away synapses and lower for more-active neurons. Neurons that hit their outgoing or incoming synaptic bound cannot build additional outgoing or incoming synapses, respectively. (c) The *Seek-and-prune* growth involves a process in which active neurons seek nearby neurons to synapse with. Synapses compete with each other through a “permanance” factor that depends on presynaptic activity and distance. Synapses with higher permance are pruned with smaller probability than those with weaker permanence (see main text and Methods for details). (b,d) Both models result in the formation of hierarchical, spatially concentrically arranged, and retinotopicaly organized visual areas.

We call the first alternative *race-to-threshold* growth (Fig. 5a-b). This rule begins with all-to-all connectivity in the form of silent synapses from LGN neurons to cortex and within the cortical sheet. The silent connections accumulate resources ferried from the soma; once the accumulated resources exceed a threshold that depends on the soma-to-synapse distance and presynaptic activation (with a smaller threshold for higher presynaptic activity and shorter soma-to-synapse distance; the threshold shares the same mathematical term Δ*_i j_* of Eq. 1: *T_i j_* = *T Δ_i j_*), the synapses become non-silent. As the silent synapses of a neuron become non-silent, the in and out-degree of the neuron increases up to a maximal in and out degree. Once a neuron becomes in- or out-degree saturated, no further synaptic resource is available for in- and out-synapse building. As a result, a ‘race’ for synaptic resources occurs, leading to the activation of synapses with the smallest thresholds first.

The second alternative that we propose is a *seek-and-prune* growth rule (Fig. 5c-d). This model begins without any connectivity between neurons. Spontaneous activity promotes the initial stochastic formation of synapses, which are then selectively pruned based on their stability. Stability is measured by a quantity called ‘synaptic permanence’, which is higher for shorter synapse-to-soma distance and synapses formed by higher presynaptic activity. This type of selective synaptic pruning is a significant feature of the developing nervous systems and is important for rewiring mistargeted axons and consolidation [42, 64, 75, 76, 77].

The functional form of the synaptic permanence is the same as the growth rate in the competition model, Eq. 1. Both rules result in qualitatively similar architecture, structure, and topography (Fig. 5b,d) Despite their endpoint similarity, we will see later that the different rules yield distinct predictions about the developmental trajectory of visuotopic organization.

### Spatial receptive field structure

Primate visual neurons have smaller foveal receptive fields with increasingly larger peripheral receptive fields within and across areas [78, 70, 79, 80, 81] — the emergent hierarchical receptive field structure of our model shows the same signatures, Fig. 6. Specifically, the log-conformal mapping from the eye to LGN also induces an eccentricity dependence on LGN field sizes, with larger fields for more eccentric retinal locations. Each cortical area inherits this qualitative eccentricity dependence, Fig. 6a (the eccentricity dependence is inherited, not amplified, by the cortex, SI Fig. S3). We further observe in Fig. 6a that at very large eccentricities near the edge of the visual field (which has not been examined closely in experimental work), the model predicts a mild downward trend of visual receptive field sizes in each area. This finding holds across varied network sizes and across the different growth rules as well as by direct implementation of the GWM-S process.

**Figure 6:**
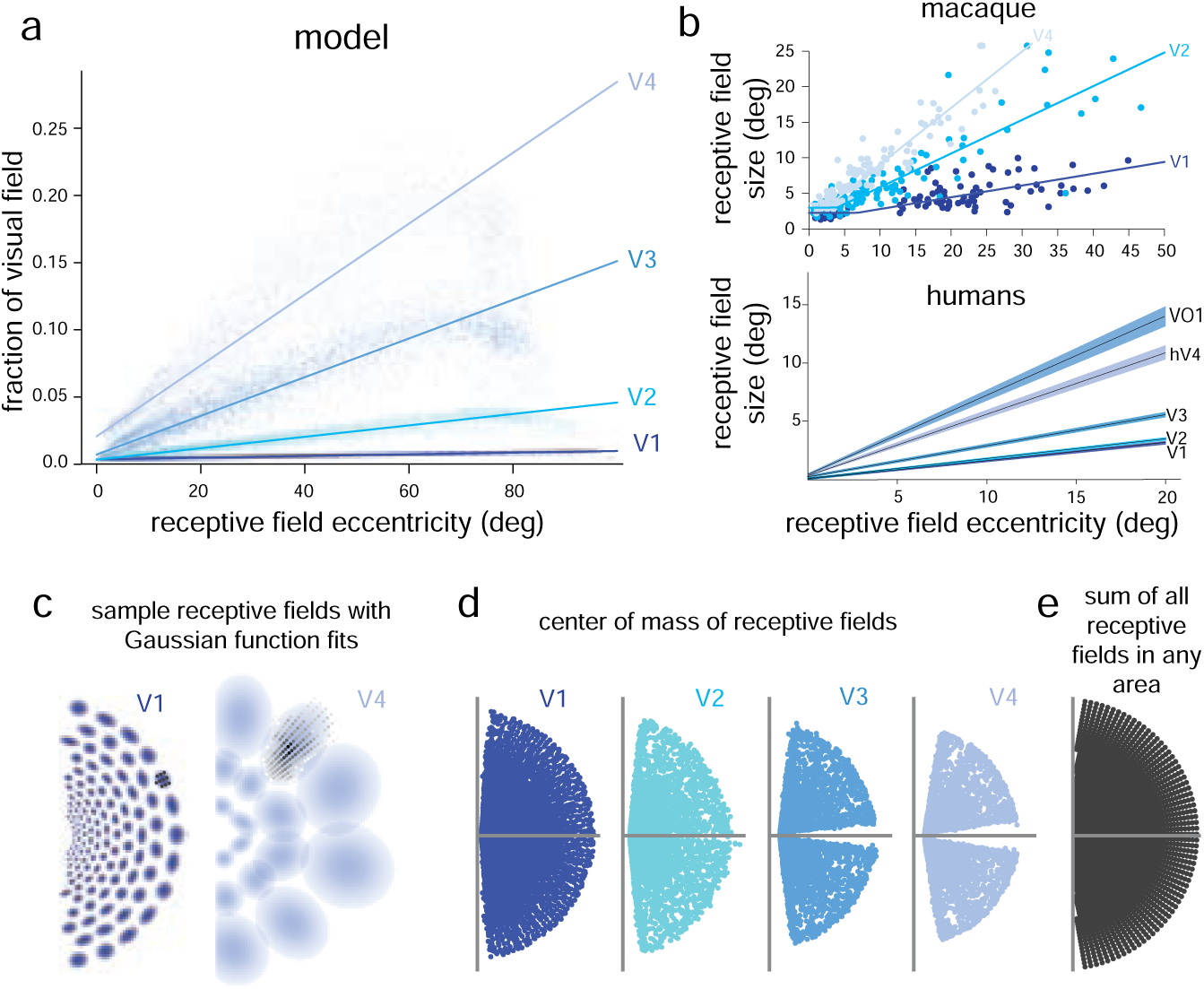
Receptive field structure in formed network. (a) Receptive fields grow with eccentricity in each visual area, with a small but significant dip at the most eccentric locations, and grow along the hierarchy. (b) Experimental data showing the eccentricity dependence of visual receptive field size from macaque[78](top) and from humans[79](bottom). (c) Example receptive fields from area V1 and V4 in the model. (d) The center of mass of receptive fields appears to show an apparent repulsion away from the horizontal and vertical meridians (shown in gray), with larger repulsion in higher areas. (e) Nevertheless, receptive fields for any area completely sample the entire visual field.

Next, the receptive fields increase in size going up the hierarchy, Fig. 6a, also qualitatively consistent with results in macaques and humans, Fig. 6b [78, 70, 79, 80, 81]. Sample receptive fields in V1 and V4 can be seen in Fig. 6c. The ratio of receptive field sizes between successive hierarchical areas should approximate a geometric progression (see next section for the reason why), but this does not hold precisely because of details like the finite cortical sheet, its geometry, and the non-isotropic shapes of local connectivity. The macaque and human data also do not exhibit a simple or consistent pattern of size progression, Fig. 6b.

Additionally, when we plot the receptive field centers for all neurons in an area, the vertical and horizontal visual meridians appear underrepresented, suggesting a repulsion of the field centers away from the meridians, 6d. On investigation, we find that this is an artifact of the fact that the horizontal and vertical visual meridians correspond to cortical area boundaries and crossing a meridian corresponds to a representational switch to the opposite cortical hemifield, with truncated receptive fields near the boundary. Defining the center-of-mass of a receptive field as the field center will result in an apparent repulsion of the field center away from boundaries, here and likely also in experiments. When the receptive fields of all neurons within an area are summed, they span all parts of the visual field uniformly, contrary to the apparent underrepresentation of the vertical or horizontal medians when using only field centers, Fig. 6e).

Finally, we observed another interesting emergent model property: Define a cortico-visual field as the patch in the visual field that a cortical cell responds to, and define a cortico-LGN field as the patch of LGN that a cortical cell responds to. The growth model generates distortions – in the form of elliptical deviations from circularity – in its cortico-LGN fields, in a way that generates circular cortico-visual fields, SI Fig. S4 [82]. Had the cortico-LGN fields been circular, the cortico-visual fields would have been more elliptical. Thus, the growth process seems to perform an automatic “whitening” of the shape of the cortico-visual receptive field, an experimental prediction.

### Connectivity

Unlike previous approaches to modeling visual cortex topography based on self-organizing maps [83], the present approach results in a complete neural circuit, with granularly specified synaptic connectivity between neurons. As already seen, it possesses the gross architecture (multiple discrete hierarchically connected areas), spatial layout (concentrically arranged areas), and topography (retinotopy and polar angle reversals) characteristic of primate visual systems. Further, the model generates a detailed connectome that can be analyzed to predict synaptic connectivity across visual cortical areas.

#### Sparse, hierarchical, and convolution-like connectivity in retinotopic coordinates

We target individual cells in the formed network, tracing their downstream and upstream partners across areas, in a dense computational version of anterograde and retrograde tracing [84] or connectomics [85], Fig. 7. A cell in a given hierarchical area sends outputs to a sparse subset of cells in the downstream area, and receives inputs from a sparse set of cells in upstream areas, Fig. 7a. This sparse set of inputs and outputs is spatially clustered in the cortical sheet (Fig. 7a). When organized based on retinotopic locations, these inputs and outputs form regularly-shaped local patches (Fig. 7b). This connectivity closely resembles the tiled local spatial organization posited to exist in brains [86, 87, 88, 89], in which neurons in each area pool inputs from the previous area through a similar pooling function (a convolution-like kernel), and which is built by hand into models that provide the best quantitative match to date to electrophysiological data from several visual areas and to behavioral data in primates and rodents [90] (Fig. 7a, far right).

**Figure 7:**
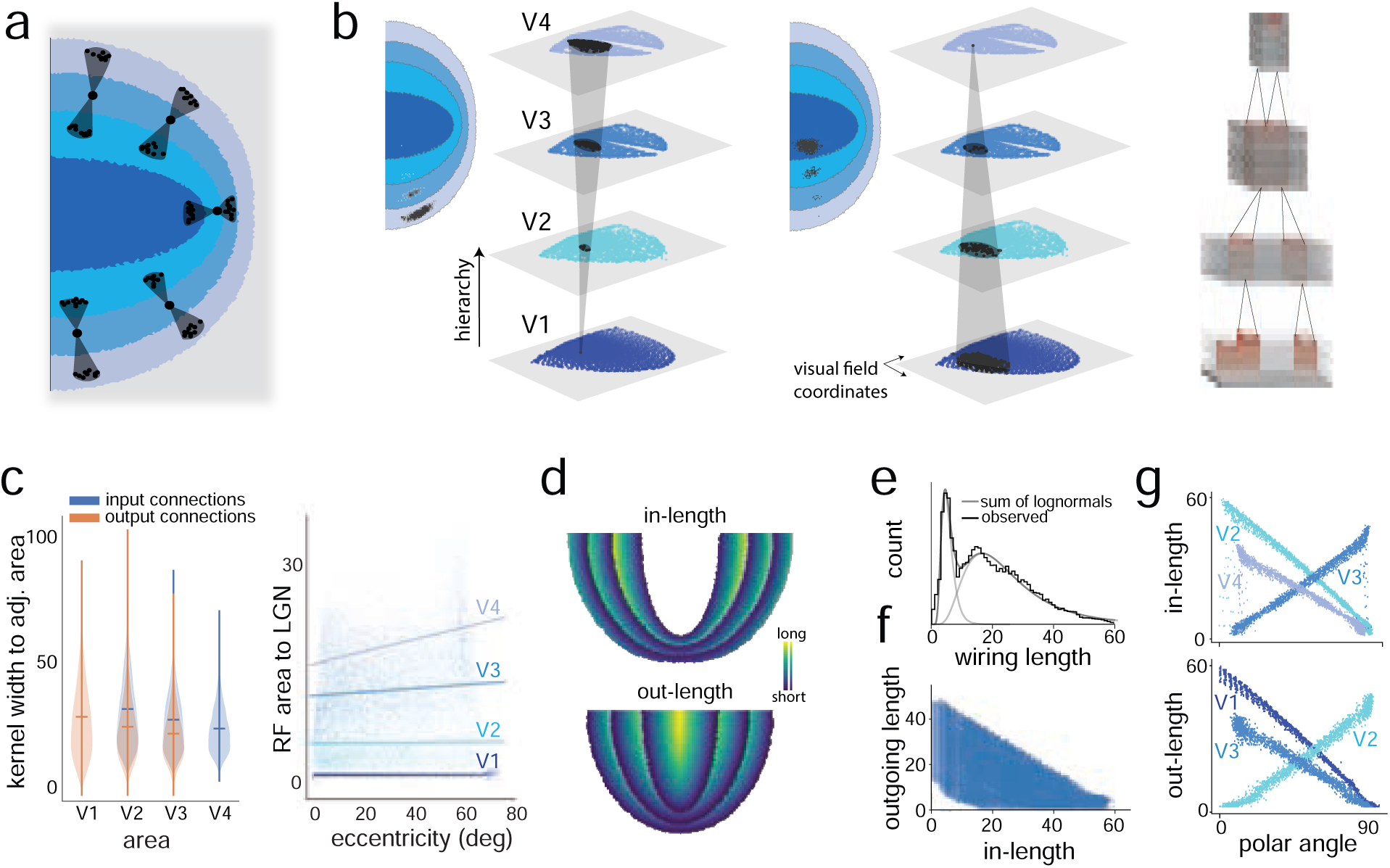
Connectivity in the grown network. (a) Computational connectomics at 5 neurons shows the incoming and outgoing connections are both sparse and local, similar to a local filter motif. (b) Left: A ‘tree’ of outgoing connections showing all neurons in the grown visual cortex that receive information from a given neuron in V1. Right: a similar tree of incoming connections, showing all neurons in the visual cortex that send information to a given V4 neuron. (c) Violin plots showing the distribution of the size of the connectivity patch on adjacent areas through input (blue) and output (orange) connections to neurons in a given area (d) A map of in-connectivity lengths and out-connectivity lengths in the formed visual cortex. Although V1 received inputs from LGN, the connectivity lengths of those synapses have not been shown here. (e) Empirically observed distribution of connectivity lengths across the entire network. (f) In-connection lengths are inversely correlated with out-connection lengths, shown for all neurons with both in and out connections in the cortical sheet (i.e., V2 and V3). (g) For large eccentricities (here chosen as larger than one-third the extent of the visual field), the magnitude of polar angle is correlated with the in-connectivity length of V3 neurons and the connectivity length of V2 neurons; it is also inversely correlated with the in-connectivity length of V2 and V4 neurons and the out-connectivity length of V1 and V3 neurons. In all cases the network used has been constructed through direct implementation of the GWM-S process; similar results are observed when explicitly using either of the three proposed biologically plausible growth rules (see SI for details).

The convolution-like connectivity profile widths are roughly similar across areas in the model (Fig. 7c left). Thus, the increase in receptive field size in higher areas of the model results from the hierarchical nesting of similar-sized local spatial kernels rather than from an increase in the kernel width between areas. As a result, the expected scaling of receptive field sizes across areas (the ratio of receptive field sizes between adjacent levels in the hierarchy) is a geometric progression. However, if spatial gradients in the cortical sheet were to modulate the maximum synaptic weights per neuron, they could affect field sizes beyond the effects included in this simplest version of a growth model. Similarly, variations of cortical connectivity width with eccentricity are weak, so that the systematic increase in receptive field size with visual eccentricity is largely inherited from the input projections to cortex, (Fig. 7c right).

#### Wiring length and location on cortical sheet

We next consider whether there are systematic variations in connectivity length within and across formed areas. Coloring cells on the cortical sheet based on the length of the input wiring reveals a sawtooth-like in-connectivity length profile, in which neurons near the inner edge of each area receive the shortest connections and those at the outer edge receive the longest connections, Fig. 7d (top), as expected from Figure 4a. Conversely, for outgoing connections, neurons at the outer edge of each area send the shortest connections, while those at the inner edge send the longest ones, Fig. 7f (bottom). Pooling together all synapses from all neurons in V1, V2, V3 and V4, we find a bimodal distribution of wiring lengths that is well-fit by a sum of two lognormals, Fig. 7e.

Neurons with long out-connections have short in-connections, and vice versa, resulting in an inverse correlation between in-connectivity and out-connectivity lengths, Fig. 7f. This pattern suggests a roughly constant summed in- and out-wiring length across neurons in an area. Finally, the relationship between wiring in-length and polar angle flips when moving along the hierarchy: in-connectivity length is negatively, then positively, then negatively correlated with polar angle for V2, V3, and V4, respectively (Fig. 4g, top). The out-length-polar angle correlations follow a similar pattern (Fig. 4g, bottom).

Altogether, the connectome of the grown visual system model reveals how a hierarchically stacked convolution-like architecture may be embedded into a locally-connected and spatially flat cortical surface with mirror-reversing retinotopic maps. This is the first model of the primate visual hierarchy to generate connectivity de novo; thus, these constitute a rich set of novel predictions for the next generation of connectomics work following the first efforts undertaken in the visual pathway [85].

#### Trajectories of retinotopy development

Across all growth rules, we already noted a similar developmental endpoint; we also observe a key common characteristic of the trajectory to the endpoint — the development of retinotopy in the visual cortical hierarchy is sequential, with retinotopy emerging and stabilizing in lower visual areas before higher areas. This constitutes a robust and invariant prediction of all the growth models. To some extent, the maturation dynamics of the visual pathway will depend on the spatiotemporal properties of the retinal waves that drive it. But under the assumption of uniformly distributed waves (such that all parts of the retina are activated with equal probability on the time-scale of development), the qualitative properties of emergence are relatively independent of the detailed form of the retinal waves: cortical hierarchy, spatial physical organization, and topography with reversals emerge from the growth rules, regardless of the details or source of the spontaneous activity that drives the process. This relative independence on the one hand reaffirms the relationship of our models to the GWM-S process. And on the other hand, it also helps explain the experimentally observed invariance of sensory hierarchy development in cortex even in animals born without a retina or with early retinal damage [91, 92], assuming that spontaneous activity is present in some downstream area. Our model therefore provides the strong prediction for future work in development that spontaneous activity covering the visual space on average, regardless of the detailed statistics of the activity, should be sufficient for architectural hierarchy, spatial organization of areas relative to each other, and mirror reversals across area boundaries. Our model suggests that finer spatiotemporal characteristics of the spontaneous activity are probably important for more detailed structure emergence and further refinement.

#### Discriminating between growth models

We next ask whether, despite their commonalities, the learning rules generate transient connectivities that are distinguishable during development. We examine retinotopy development at the intermediate period after V1 and V2 formation and while V3 is partially organized (Fig.8). Under the heterosynaptic growth rule, all cortical neurons beyond V1 and V2 are connected and receive input drive. A rough retinotopy is rapidly established across all neurons in proto-V3 (Fig 8b, light blue curve), then the retinotopic organization is gradually sharpened *in situ* (Fig 8b, darker blue curves).

**Figure 8:**
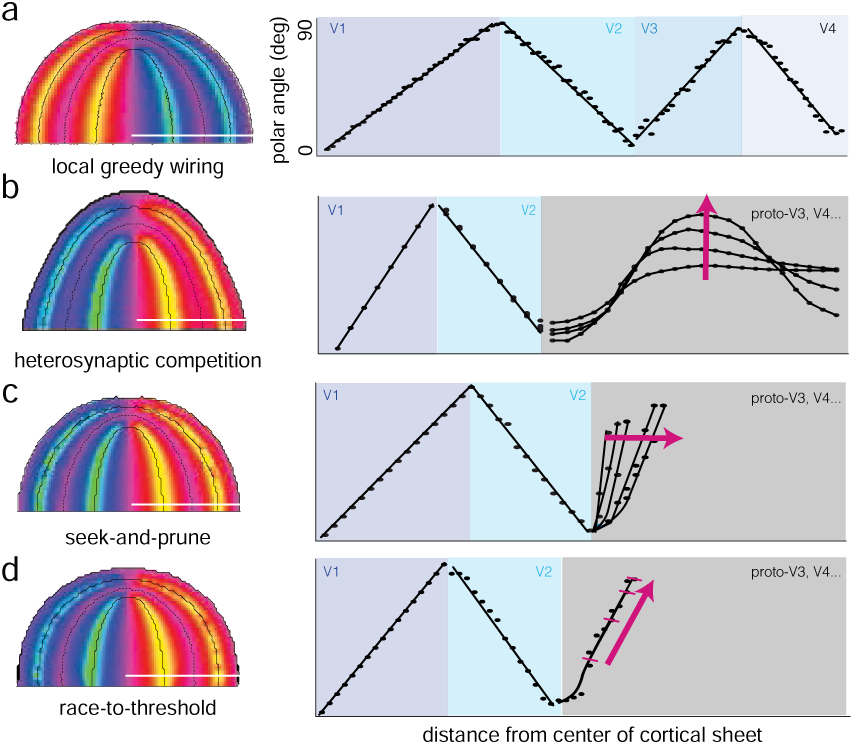
Distinguishable developmental trajectories under different growth rules. (a) Mature cortical sheet under the GWM-S process (left). Horizontal white line: Iso-eccentricity slice along which the polar angle varies as shown (right). (b-d) Layout of mature cortical sheet for three wiring rules (left column), and quantification of the development of polar angle tuning (right column). Developmental trajectory around an intermediate stage of growth after the formation of V1 and V2 but before the complete development of retinotopy in V3. Magenta arrow shows the progression of developmental stages.

By contrast, under the seek-and-prune and race-to-threshold growth rules, neurons beyond V1 and V2 exhibit no retinotopy initially, Fig. 8c-d. Under both these rules, the retinotopically structured part of proto-V3 slowly expands outward. Under seek-and-prune growth, most neurons in proto-V3 are unconnected to V1 or V2, and neurons in the region slowly gain connectivity to V2 inputs, forming an area V3. However, at any point, the partially grown V3 contains a complete map of the visual field (i.e., coverage of all polar angles and eccentricities), Fig. 8c. As V3 expands, the polar angle map stretches out in tandem. Under race-to-threshold growth, neurons in proto-V3 represent only a small range of polar angle (the angles most similar to those at the V2 edge). However, the angles represented by these proto-V3 neurons remains similar to the angles these neurons will represent in mature V3. Over time, neurons further away in the proto-V3 gain a polar angle representation, steadily covering a larger range of angles over time in tandem with the gain in polar tuning across proto-V3 (Fig. 8d).

Across models, the shorter within-area connections are predicted to stabilize earlier than the longer within-area connections. In sum, we have explored a set of biological growth rules that all follow the local GWM-S principle, but differ in some biologically important ways that we predict can be distinguished based on specified and measurable differences in their developmental trajectories.

### Generalization to other cell types: Interneuron emergence

Only a small number of parameters modulate the growth dynamics, across all rules. The distance scale *d*_0_ in Eq. 1 and the activity propagation attenuation factor γ in equation 3 are critical in shaping outcomes and are common across rules. Two additional shared parameters set the maximum in-degree and out-degree of each neuron — in race-to-threshold and seek-and-prune, these are *k^in^*, *k^out^*; in heterosynaptic competition, these are (the ratio of) the maximum per-synapse and per-neuron weights *w*_max_, *W*_max_. These degree parameters affect the relative sizes of successive areas, SI Fig. S2, but do not qualitatively change the architecture. Finally, each growth role requires growth rate parameters which affect numerical stability and convergence, but which do not qualitatively alter the final network; for heterosynaptic competition, these are set by η (growth rate),ε (competition strength), and ε*_w_* (activity propagation threshold); for growth-to-threshold, these are set by η (growth rate) and *r*_0_ (normalization of accumulated resources); for seek-and-prune, these are *n_draw_* (number of synapses neurons attempt to form in each growth iteration). Combined with structural specification of the cortical sheet and LGN geometry, which are held fixed across rules, this is the exhaustive list of parameters.

For the two critical parameters related to activity propagation γ and distance scale *d*_0_, the growth dynamics are robust to variations (up to 5 orders of magnitude of variation in these parameters, cf. SI Fig. S5), However, we found that there is a boundary in the γ, *d*_0_ space across which the network switches from developing a hierarchical architecture to instead developing a non-hierarchical one with purely local connectivity everywhere (Fig. S5).

The latter connectivity is reminiscent of the projection patterns of inhibitory interneurons in cortex, predicting that genetic specification in this γ, *d*_0_ space could be a developmental control knob that is set differently across cell types to govern their projection patterns.

Thus, to investigate the potential for simultaneous emergence of both projection patterns, we created an undifferentiated cortical consisting of two types of cells Fig.9a, interleaved in a 1:8 ratio (see *Methods* for details). Regardless of type, all neurons interact with and drive each other. They all compete together to innervate the same set of potential neural targets. Remarkably, this mixed-neuron system again forms a hierarchy of discrete areas that are concentrically organized on the cortical sheet, Fig. 9b,left.

**Figure 9:**
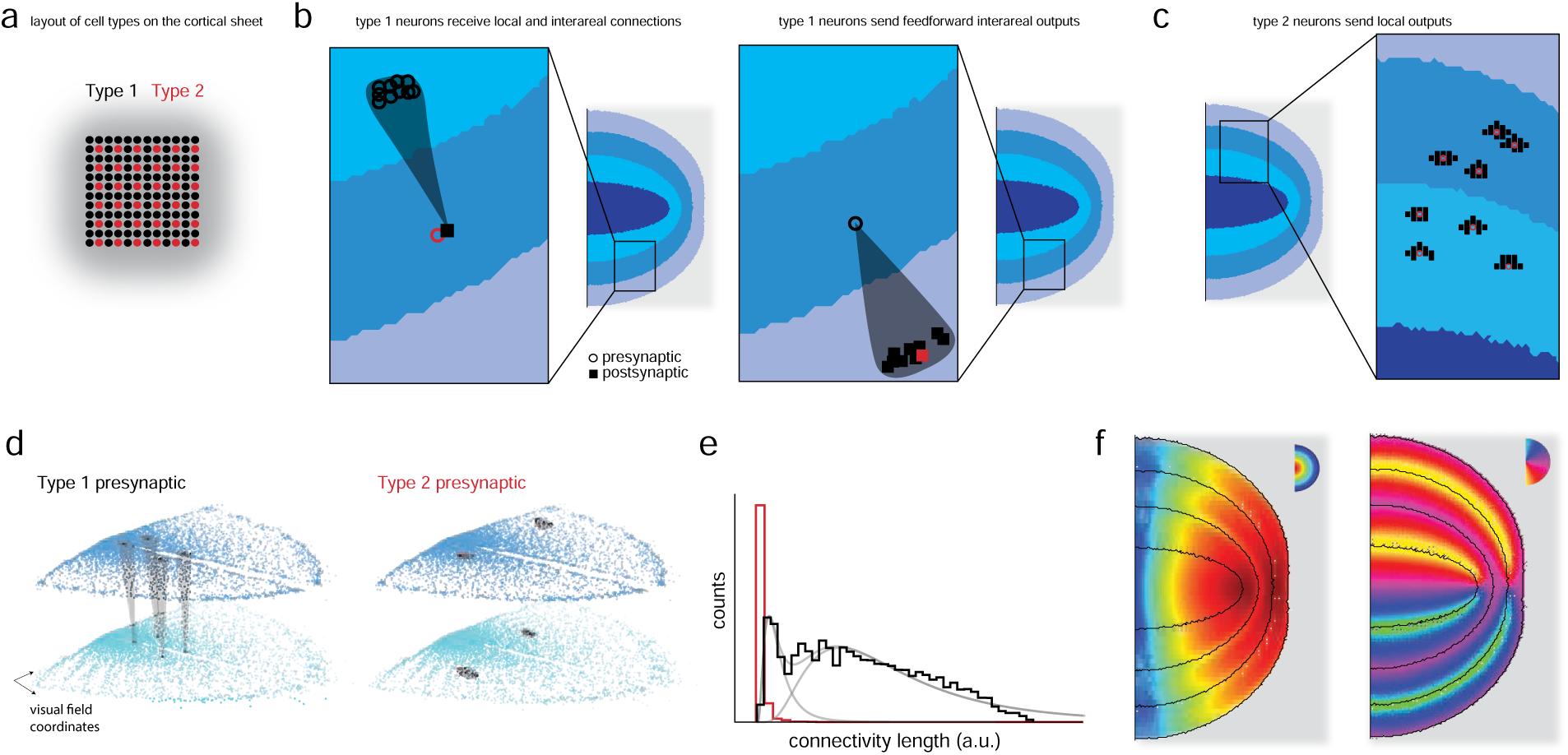
Incorporation of cell types and emergent cell-type dependent connectivity motifs from growth: (a) A schematic of layout of the 2 cell types in a square lattice on the cortical sheet (b) Simulation with the incorporation of the two cell types continues to result in the formation of concentrically arranged areas. Within these areas, cells of type 2 form local connections (open circles designate pre-synaptic partners; solid squares designate post-synaptic partners) (b), whereas cells of type 1 receive both local and feedforward connections (c,*left*), but form only feedforward connections (c,*right*). (d) Arranging the neurons according to their hierarchy and receptive field coordinates (similar to Fig. 7(a-b)) similarly shows that cells of type 1 project feedforward into the next hierarchical area, whereas cells of type 2 project only locally to neurons in the same area. (e) Histogram of all wiring lengths, with output connections from cell type 2 neurons shown in red, and output connections from cell type 1 neurons shown in black. (f) The associated eccentricity (left) and polar angle (right) maps.

Connectomic tracing of the in- and out-projections reveals that type 1 cells exhibit local spatial projection patterns to successive areas in the hierarchy, Fig. 9b, like the single-cell-type results. We will call these projection neurons. Type 2 cells exhibit purely local within-area projections, Fig. 9c. We will call these interneurons. Interneurons, like projection neurons, receive both local inputs and feedforward inputs from the preceding area. The differences in the connectivity of the two cell types can also be visualized when neurons are organized based on their retinal receptive field positions, Fig. 9d, and can be quantified in a wiring length histogram, Fig. 9e. As with the single cell-type results on the spatial width of neural projections as a function of level in the formed hierarchy (Fig. 7c), the interneuron connectivity width is also constant across levels of the hierarchy and determined by the out-degree of the neurons, Fig. 9c.

The formed areas exhibit qualitatively similar mirrored retinotopic maps in areas V1-V4 as before, Fig. 9f, albeit slightly noisier. This network constitutes a skeleton (connectivity graph) model of the ventral visual hierarchy with both projection neuron and interneuron connectivity patterns simultaneously formed from the same set of simple growth rules.

### Generalization to other sensory modalities: Auditory hierarchy

The robust hierarchical structures that emerge from our developmental rules suggest that other sensory modalities and sensory hierarchies in different species could be modeled in a similar way.

In the case of vision, the input topography was specified by the visual field projected onto the retina. The placement of LGN relative to cortex, in our case with LGN nearest to one edge of the cortical sheet, resulted in concentric organization of hemi-circular or horeshoe-like areas around the primary cortex. Had the LGN been aligned to the center of the cortical sheet, cortical areas would fully surround the primary area.

The radial topography and polar angle alternations were similarly a function of LGN geometry: the mirror reversal of maps across area boundaries corresponded to polar angle reversals because iso-polar-angle contours ran parallel to the LGN boundaries. If the LGN geometry were flattened so that iso-eccentricity contours ran parallel to its boundaries, this would result instead in a eccentricity map alternations along with a more-smooth mapping of polar angle SI Fig. S6. Examples with different LGN positioning relative to the cortical sheet and LGN geometry are given in Fig. 10 and SI Fig. S6. Differences in these initial geometric properties might be partly responsible for variations across cortical modalities and species.

**Figure 10:**
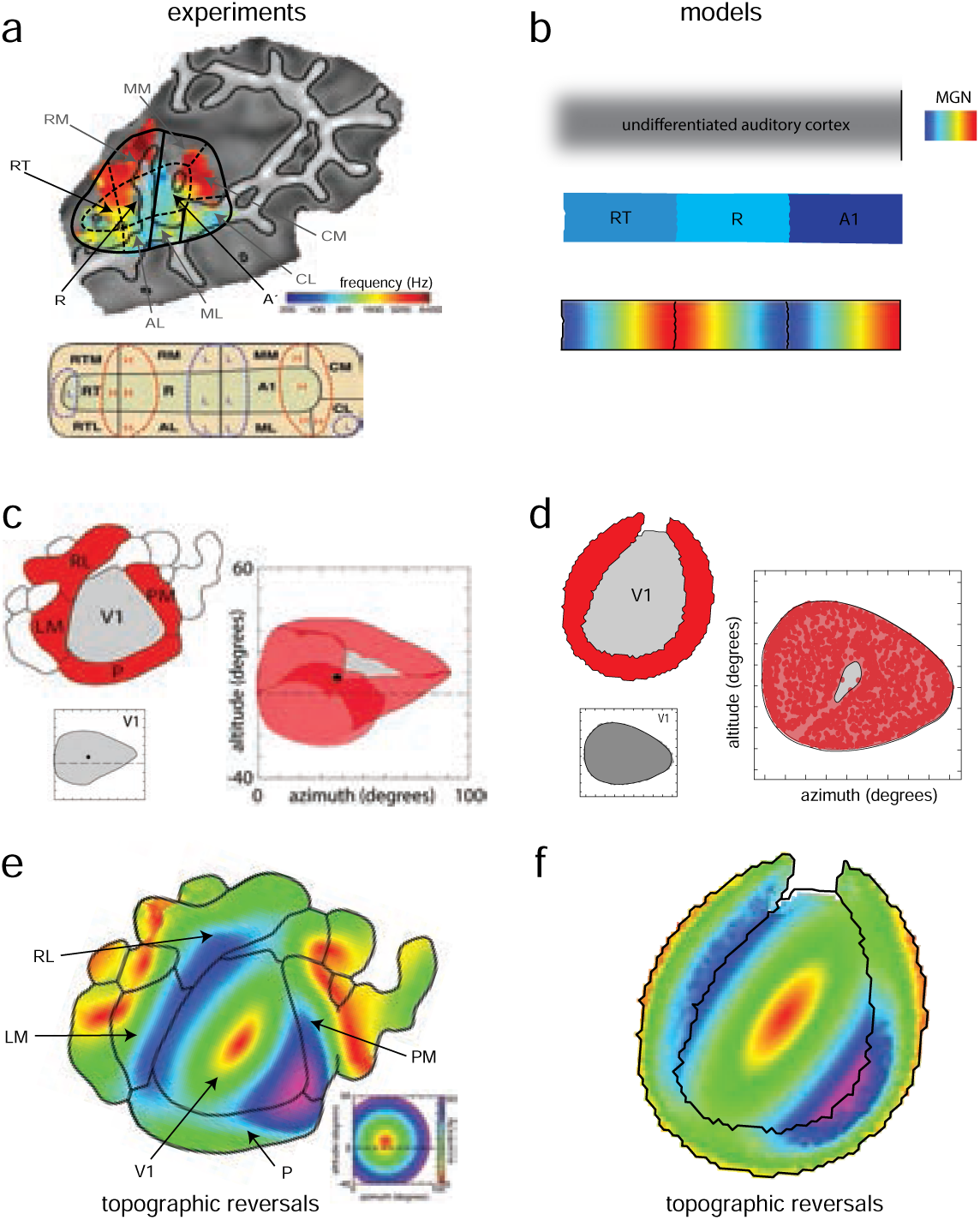
The greedy wiring minimization principle generates cortical processing hierarchies across sensory modalities and species. (a) Top: A tonotopic map of auditory cortical areas from fMRI in humans exhibits smooth gradients in auditory frequency tuning as well as mirror reversals between areas [10]. Bottom: Schematic of the auditory cortical layout (H and L mark high- and low-frequency tuning, while A1, R, and RT designate successive areas in the auditory processing hierarchy). (b) Simulation of the local GWM-S process applied to a strip of undifferentiated cortex with spontaneous input assumed to arrive from a cochlea, results in a discrete set of auditory areas with mirror reversals of tonotopy. (c) (Top left) A map of the mouse visual cortex spanning areas including V1, P, PM, LM, RL [93]. The coverage map in visual space of V1 shown in gray (bottom left) is almost entirely covered by the coverage map of the higher areas, P, PM, LM and RL (right) with a marked hole in visual space representation. (d) A simulation of the GWM-S process on a cortical sheet seeded with the appropriate geometry results in the formation of a hierarchical second area (red) that surrounds the initial putative V1 (gray). the greedy wiring simulation results in similar coverage maps of V1 (bottom left), and a similar hole visual space representation of the higher area (right). (e) A retinotopic map of the mouse visual cortex, and (f) of a network grown with the GWM-S process.

In the auditory system, the sensory organ is the cochlea. Its tapered geometry induces a one-dimensional tonotopic mapping between sound frequency and responding regions of the cochlea. Like the retinotopic LGN projection into undifferentiated visual cortex, we assume a tonotopic cochlear projection via the auditory thalamus MGN into undifferentiated auditory cortex. We model the undifferentiated auditory cortex as a rectangular strip that roughly matches the geometry of the core-belt auditory regions, Fig. 10a, with MGN projecting toward the caudal end [94, 95], Fig. 10b.

Competitive growth drives the formation of a discrete auditory hierarchy with tonotopic mirror reversals at the area boundaries. We equate the formed regions with A1, R, and RT, which, based on response latencies, are believed to form a hierarchy of processing areas in auditory cortex [10, 94, 95], Fig 10c. Given the relatively less-detailed knowledge of maps in the auditory cortex, our hope is that information flow and connectivity predictions from this model might provide interesting hypotheses about auditory cortical organization for experimental study.

### Generalization to other species: Mouse visual cortex

The murine visual cortex has a less clear hierarchical structure than primates. The primary visual area V1 projects to a number of regions, including P, LM, AL, and RL. It is a subject of debate whether these are distinct processing areas or whether together they define a second stage in a processing hierarchy. Recent work with wide-field calcium imaging generated a detailed retinotopic map of primary visual cortex and other extrastriate areas, Fig 10a (top) [93], revealing that secondary areas P, LM, AL, RL do not individually cover the visual field, but together cover most of it, apart from a puncture near the center, Fig 10b (bottom, right). We ran the local greedy wiring minimization (GWM-S) simulation with a neural sheet matching the mouse visual cortex and seeding it with an input geometry that matches the mouse primary visual cortex. The result reproduced the retinotopy and the visual field coverage of V1 and the surrounding areas, Fig 10b.

Thus, the same learning rules, applied to inputs from different modalities or to different cortical geometries for different species, generate distinct sensory hierarchies in different animals. The framework of our model thus provides detailed predictions at the neural and connectomic levels across sensory modalities and species.

## Discussion

In the present work, we show how synaptic-level subcellular learning rules and cellular-level competition can organize an unstructured cortical sheet into a set of discrete, hierarchically connected brain regions, with their topographic mapping inherited from the metrics of the low-level sensory space and from the geometry of the cortical sheet. Thus, the model parsimoniously explains how, driven by visual inputs, cortex would develop a visual processing pathway with discrete brain areas and hierarchical architecture, spatially concentric organization of these areas on the cortical sheet, and topographic structure in the form of retinotopy and polar alternations, and how. The same rules and model, driven by auditory inputs, develop an auditory processing hierarchy. The same model acting on a different cortical geometry also generates a mouse-like visual cortical processing stream. Thus, the model provides a mechanistic hypothesis for fascinating discoveries on the pluripotency of cortical development [28].

Burgeoning data from connectomics and transcriptomics, as well as neural imaging and activity data, demand models that link the disparate levels from genes and cells to connectivity, local circuit structure, and brain organization. Under the hypothesis that the small number of parameters of the growth rules are controlled by genetic guidance, our model also provides predictions for linking all the way from gene expression to complex emergent structure. These parameters control whether a cell’s connectivity is projection neuron-like or interneuron-like and the thickness or size of each discrete cortical area. Thus, the model predicts that key combinations of gene expression map onto the control knobs of the growth model to determine the projection patterns of cell types and the macroscopic features of areal organization, such as area size, and should vary in specifically defined ways according to the model to generate different cell types (projection neurons and interneurons), different sensory cortices (visual and auditory), or different species (primate and mouse).

The proposed mechanisms of structure emergence via competitive growth rules that implement local greedy wiring minimization result in self-organization driven by spontaneous activity. Thus, they contrast with the traditional chemoaffinity hypothesis of Sperry [17], in which genes instruct the detailed organization of synapses. This is consistent with the finding that even in the fly, an exemplar of chemoaffinity-guided development, spontaneous activity plays a critical role in brain development [32, 38, 30, 52, 53]. Our model provides a potential mechanism for how self-organized structure emergence could work in tandem with or in addition to detailed genetic specification. Moreover, our model goes beyond the idea that spontaneous activity drives refinement of genetically specified wiring; here, spontaneous activity also plays a role in setting gross features like discrete region formation, hierarchy emergence, and global polar angle reversals.

### Relationship to existing models: global wiring minimization and visual cortex organization

Several works in the literature focus on global wiring optimization, showing that ocular dominance and orientation preference maps can arise within the framework of optimizing a global synaptic wiring length objective between co-tuned neurons [96, 72, 54]. These works contrast from ours in two ways: They focus on algorithms for global optimization without biologically plausible processes to achieve such optimal solutions in the brain; and there is no demonstration that global wiring optimization as a principle would generate discrete brain regions, hierarchical order, spatial order, or topographic mirror reversals. Another wiring minimization approach involves supervised learning: it begins with a pre-specified architecture (discrete hierarchical brain areas) and connectivity (spatially local), then generates neural tuning and layouts within each area by maximizing performance on a visual task while minimizing total wiring length through gradient learning [74, 97]. These global optimization approaches do not lend themselves to the creation of mechanistic models of cortical development because global wiring length minimization is an NP-hard problem and because they begin with a prespecified connectivity and focus on spatial optimization based on that connectivity.

Alternatively, self-organizing map approaches have sought to model visual topographic signatures, including ocular dominance [98, 99, 100], the development of the log-conformal map in primary visual cortex[101] and more recently, multiple visual field maps in early visual cortical areas[102, 83]. These approaches generate a spatial organization based on prespecified tuning profiles rather than a prespecified connectivity, but the resulting models do not generate biologically plausible synaptic growth rules or result in circuit-level connectivity for how the assumed tuning profiles might arise.

By contrast, we cast development as driven by local growth rules fueled by spontaneous activity, with neuron-level synaptic constraints and local competition, in a process that precedes the emergence of specific visual feature tuning, consistent with experimental findings. The unfolding dynamics naturally generates a hierarchical multi-scale structure with a full neural-level connectome, in which connectivity is not globally optimized but is the result of local search process where neurons sequentially stabilize their shortest connections.

Our work can be viewed as extending [35], which explains some properties of the visual system by growing a network from developmental rules. Our model extends the quasi-one-dimensional structure of [35] into two dimensions to capture the different topographies of both retinotopic eccentricity and polar angle (smooth versus mirror reversing topographies, respectively), naturally generates inter-areal alternations (in contrast to [35], which requires that the degree of saturation in subsets of neurons is specifically modulated in time to generate alternations), and generates a hierarchical organization (in contrast to [35], in which all resulting connections are from V1 to other areas rather than a hierarchy).

### Predictions

The model makes numerous predictions. Several include mechanisms to generate previously known aspects of structure (hierarchy, spatial organization, and topography) in a highly compact and parsimonious way, while predicting that region boundaries defined by two independent measures – synaptic hops or hierarchy index, and topographic mirror reversals – should coincide. In a real sense, because the model includes both a growth process and a full final connectome, it also generates an uncountable number of novel predictions, as development can be simulated and the outcomes predicted for any cortical geometry and topology with or without cortical lesions or retinal scotomas, any input geometry, and different settings of the few parameters of the growth rule. We have described several specific predictions along the way in the Results and in the Discussion above; we list a few of them here as well: 1) spontaneous activity that covers the visual space on average, regardless of the detailed statistics of the activity, should be sufficient for the emergence of architectural hierarchy, the spatial organization of areas relative to each other, and mirror reversals across area boundaries; 2) More-detailed spatiotemporal patterns of the spontaneous activity are important only for further refinement and more-detailed structural and functional characteristics. 3) Dip in field size with eccentricity at large eccentricites; 4) Sequential development of retinotopy up the hierarchy, such that retinotopy emerges and stabilizes in lower visual areas before higher visual areas, true across all the rules; 5) More-circular cortico-visual receptive fields compared to cortico-LGN fields; 6) Two critical factors of the growth process and neural/synaptic saturation determine the width and size of each discrete formed region when cortical size is held fixed; 7) Hypothesized relationship between growth process parameters and genetic control is predicted to enable simple scaling of the size of each cortical are by simple genetic manipulation; 8) Different synaptic growth rules, all under the greedy wiring minimization principle, have developmental trajectories that differ in specific predicted ways; 9) Detailed predicted connectome for the auditory processing pathway in primate cortex and for visual pathways in mice [103, 104] to assist in mapping out these less-well characterized hierarchies; 10) Saw-tooth pattern of wiring length when moving from the center of V1 radially outwards in cortex across areas, leading to a systematic (alternating inverse and direct) relationship between polar angle and wiring length across areas; 11) Inverse relationship between pre-synaptic wiring length and post-synpatic wiring length for feed-forward connections between areas; 12) Bimodal distribution of wiring lengths for feed-forward connections between areas; 13) First full primate visual pathway connectomic model including the effects of the log-normal magnification of the fovea, constituting a rich set of additional predictions for the next generation of connectomics work in the visual pathway [85, 103].

### Limitations and future work

We have focused here on articulating a compact set of rules that drive modular hierarchical structure emergence. Our model should be viewed as setting the backbone or connectivity structure of the ventral visual pathway in the late embryonic or post birth but pre eye-opening stages. Future work will focus on how the spatiotemporal structure of retinal waves could itself direct the formation of our assumed retinotopy in the LGN projection, and the emergence of some aspects of image-feature tuning, including orientation selectivity and ocular dominance columns, which occur at a later stage of development in the visual system [105].Two retinas will be required to explore the formation of ocular dominance stripes, while patterned visual inputs from natural images might be required to learn finer topographic features including orientation maps.

Our work does not yet extend to the laminar structure and specializations of individual cortical areas. The result that dialing the growth rule parameters generates an interneuron-like cell class, and that these cells become integrated into the feedforward backbone with the same growth rules, opens the prospect that it might be possible to similarly extend the present model to incorporate more distinct cell types and for a combination of self-organization and genetic specification to drive laminar organization [106]. Specifically, adding emergent dynamics in setting the growth parameters themselves could allow for rich developmental outcomes including laminar organization and different spreads in lateral connected at different hierarchical processing levels. Our model lays the groundwork to begin modeling these finer features of cortical organization. It will also be interesting to obtain extensions that generate top-down connections [107, 108]. Finally, this model can be extended to study how regional neural waves in other sensory peripheries and higher-level areas such as thalamus shape downstream connectivity [109, 110]. It will be interesting to investigate how to combine processes of parallel or simultaneous modular circuit development [14, 35] with the present sequential development model.

### Broader implications for brain organization

The present growth models or small modifications of them, within the GWM-S principle, might play a role in explaining brain organization beyond the early sensory hierarchies. Specifically, mirror-symmetric map layouts or topographic organization matching that of sensory pathways have been reported in higher brain areas [111], such as in lateral prefrontal cortex [112] the intra-parietal sulcus [113, 70], frontal areas [70], cerebellum [114], striatum [115], and hippocampus [116]. It is possible that the hierarchical map formation process with topographic mirror reversals in sensory areas could continue in some form up to higher areas in the brain, forming a sensory-coordinate layout of cognitive brain regions [112, 115].

Finally, the present model uses precisely the same heterosynaptic competition-based learning rule that was hypothesized to generate robust sequence-formation in songbird premotor circuits [46]. It is striking that the same rule, when driven with retinal inputs and a spatial embedding robustly generates a discrete, modular sensory hierarchy with various types of structure, while when driven with random inputs, and a strengthening term that is asymmetrically spike-time-dependent rather than spatially dependent, robustly generates sets of long spatiotemporal activity sequences. This striking potency of the competitive growth rule, in addition to explaining the pluripotency of sensory cortical hierarchies, is another indication that it might be a universal driver of developmental brain organization.

This hypothesis and computational model is consistent with a long history of results from developmental neuroscience in which competitive interactions and winner-take-all-dynamics lead to self-organization and structure emergence through spontaneous symmetry breaking [13, 117, 42, 43, 44, 45, 118, 119]. Neural-level competitive synaptic interactions also occur beyond development, under conditions of resource scarcity, for instance set by the rate of protein production [120], and can give rise to phenomena like synaptic clustering on dendritic arbors [121] and recovery from lesions [122, 123].

## Methods

### Cortical geometry, retinal waves, initial connectivity

In this section, we provide mathematical and implementation details common to all growth models described in the paper. In each growth-rule case, the LGN-cortical synapses and cortico-cortical synapses share a common rule, with identical parameters.

#### Geometry

Numerical simulations are initialized with two sets of neurons: 1) LGN cells, which are assumed to be driven by the retina via a log-conformal mapping, and 2) cells in the initially undifferentiated cortical sheet. LGN neurons are arranged according to the log-conformal map from a visual hemifield, a shape qualitatively similar to a semi-ellipse, which we use as our LGN geometry. The cortical sheet is modeled as a semi-infinite sheet of neurons. The straight edge of the LGN semi-ellipse is aligned with the straight edge of the semi-infinite sheet, lying above this sheet. The distance between two cortical neurons is a two-dimensional Euclidean distance on the cortical sheet, and the distance between an LGN neuron and a cortical neuron is the simple three-dimensional Euclidean distance between points on the two-dimensional Euclidean semi-sheet and the two-dimensional LGN semi-ellipse.

We assume that the LGN neurons, which receive retinal input, all start and remain “in-degree saturated”, i.e., the LGN neurons cannot accept any additional inputs from other LGN neurons or cortical neurons.

We denote the spatial position of neuron *i* by **x***_i_*. Directed connections from neuron *j* to neuron *i* are represented by the connectivity matrix *C_i j_*, whose binary elements (0, 1) represent the absence or presence of a directed connection.

Synaptic weights at the connection *C_i j_* are represented by *W_i j_*. Note that *C_i j_* = 0 implies *W_i j_* = 0, however *W_i j_* = 0 does not imply *C_i j_* = 0, as in the case of silent synapses.

#### Activity

Activity in our network model is driven by retinal waves, and this activity propagates though the LGN and cortex via the connectivity and weights formed by the growth rules. In turn, the growth rules are fueled by this propagated activity. Here we describe activity in the network given the connectivity, and then below describe the growth rules given activity.

Our activity model is highly simplified to capture a core set of properties of activity generation in the retina and propagation across the cortex. Retinal waves are spatiotemporally propagating fronts of activation. Each wave is initiated locally and spreads to cover part of the retina. Several such waves together cover the whole retina. Suppose a set of retinal waves together covers the whole retina at some timescale τ*_sa_*. Suppose that the process of cortical development and plasticity has a timescale τ*_pl_*; we discretize time into steps on this timescale, and index these times as *{t*, *t* + 1, *t* + 2, *· · · }*). We assume, based on empirical results, that these timesteps are significantly longer than τ*_sa_*. Thus, we can take retinal activation within one such time-step to be uniform across the whole retina. Mathematically, we therefore set retinal activity as *a^R^*(*i*; *t*) = 1 across all retinal positions *i* and development timesteps *t*.

We assume a fixed feedforward 1-1 connectivity between retinal locations and a topographic set of LGN neurons. Thus, *a*(*i*; *t*) = *f* (*a^R^*(*i*, *t*)) for all *i ∈ LGN* (that is, for all LGN neurons), and for all *t*. Based on empirical results we also assume that the dynamics of activity propagation from the retina into LGN and the cortex is at a timescale τ*_sp_* that is rapid relative to both τ*_sa_* and τ*_pl_*. We call these discrete time-steps δ*t* (with δ*t* 1). Thus, activity propagation in the LGN and cortex in the developmental time interval [*t*, *t* + 1] can be written by the iteration:

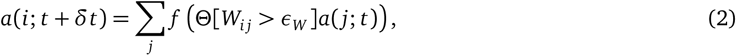

where *Θ*(*x*) is the Heaviside-theta function (*Θ*(*x*) = 0 if *x* ≤ 0, and *Θ*(*x*) = 1 for *x* > 0); ε*_W_* 1 is a small threshold on all synaptic weights, applied to prevent activity propagation across synapses with negligible strength. The iteration should be performed *N* = τ*_sa_* /τ*_s_*p≫1 times, until the next step. For simplicity, we set the neural transformation to be linear with a prefactor close to 1, *f* (*x*) = γ*x* (γ ≤ 1). With this simplification, we can explicitly sum the iteration over *N* synaptic propagation steps to obtain that at *t* + 1, activity is given by:

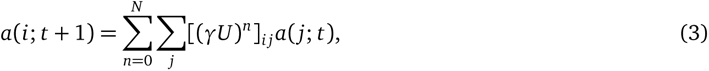

where *U Θ*(*W* > ε*_W_*) is the thresholded weight matrix.

For numerical simulations, it suffices for *N* to be larger than or equal to the number of layers desired at the end of the simulation; we choose *N* to be 5 in all of our numerical results across all growth rules. The activity reduction factor γ < 1 promotes the formation of a hierarchy by incentivizing feed-forward connections from lower layers (which will have stronger pre-synaptic activity) over lateral connections in higher layers (with weaker pre-synaptic activity).

### Heterosynaptic competition rule and implementation

In this growth rule, we assume a fully connected initial network (i.e., *C_i j_* = 1), but the weights *W_i j_* are zero, akin to silent synapses. At each time-step, weights grow at a rate proportional to Δ*_i j_*,

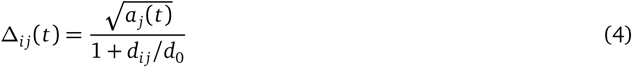

with *d_i j_* = *|***x***_i_ −* **x** *_j_|*. In other words, the rate of growth of a synaptic weight increases with the presynaptic firing rate *a_j_*, and is inversely proportional to the distance between the pair of neurons.

Within each neuron, we assume a competitive dynamics for synaptic strengthening, such that all incoming weights to a cell are decremented by the same amount when their sum hits a bound 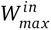, and all outgoing weights from a cell are decremented by the same amount when their sum hits a bound 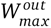. Thus, the total weight update is given by:

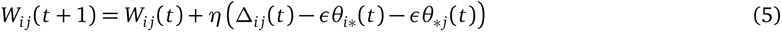

where 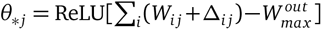 and 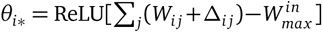. At each step, weights are clipped to the range [0, *w_max_*].

### Race to threshold rule and implementation

In this model, as above, we assume a fully connected initial network (i.e., *C_i j_* = 1) with silent synapses (*W_i j_*(0) = 0). At all future times, synaptic weights are binary (*W_i j_*(*t*) *∈ {*0, 1*}*).

Let *r_i j_* represent the accumulation of synaptic resources that are being ferried from the soma to each connection. Once this quantity exceeds a threshold that is set in an activity- and distance-dependent way, the synaptic weight becomes non-zero. Specifically,

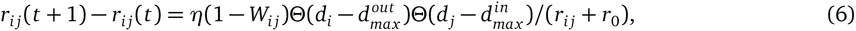

where the rate of resource ferrying is constant to all silent synapses for which the pre- and post-synaptic neurons are not out- or in-degree saturated (the out- and in-degrees are smaller than *d_max_*). For non-silent synapses, the rate of resource ferrying saturates as a function of synaptic strength (*W_i j_*). *r*_0_ is a small regularization constant for numerical stability, and does not affect the qualitative nature of the dynamics beyond this effect.

Synapse *i j* gains a non-zero weight once it accumulates resources that exceed a threshold:

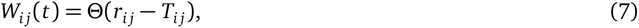

where the threshold *T_i j_* grows with the distance between neurons *i*, *j* and decreases with the level of presynaptic activity: 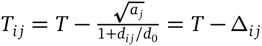. Note that the threshold here is closely related to the key quantity Δ*_i j_* from the heterosynaptic competition rule.

### Seeking and pruning rule and implementation

The two synaptic growth rules above are deterministic functions of neural activity. In contrast, seeking and pruning is a stochastic rule, where synapses are discretely and stochastically formed, then pruned. In this model, there are initially no connections, i.e., *C_i j_* = 0. Weights at all future times are binary (*W_i j_*(*t*) *∈ {*0, 1*}*).

At each time-point, we randomly select *n_draw_* neurons which are not yet out-degree saturated with probability proportional to the magnitude of their activity (initially this *n_draw_* set resides entirely within LGN neurons, since only they are active due to retinal input). Each of the *n_draw_* neurons attempts to generate *d^out^* new synapses with the set of *d^out^* spatially closest neurons to it on the neural sheet. For any given synapse, say from neuron *i* to neuron *j*, this attempt is successful if either of the following hold: (a) neuron *j* has fewer than *d^in^* synapses (it is not in-degree saturated, defined in the general setup earlier), or (b) the “permanence” *P_i j_* (defined below) of the new synapse is larger than the permanence of at least one of the existing synapses onto neuron *j*. In case (b), the synapse to neuron *j* with the weakest permanence is pruned, and the synapse from neuron *i* to neuron *j* takes its place. If neither (a) nor (b) hold, the synapse from *i* to *j* cannot be formed and neuron *i seeks* out the next-nearest neuron. This proceeds until neuron *i* (and each of the *n_draw_* neurons) successfully connects to *d^out^*.

The synaptic “permanence” of each attempted and formed synapse is given by 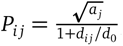, which determines how resistant it is to pruning (permanence is closely related to a key quantity used in the other rules, *P_i j_* = Δ*_i j_*).

The permanence of a synapse varies proportionally with the activity in the presynaptic neuron and inversely with the spatial distance between the pre- and post-synaptic neurons on the neural sheet.

### Parameter choices

| Biophysical growth models parameters |  |  |  |  |  |
| --- | --- | --- | --- | --- | --- |
| Seeking and Pruning model |  | Race for resources model |  | Heterosynaptic competition model |  |
| $N_x$ | 75 | $N_x$ | 75 | $N_x$ | 75 |
| $N_y$ | 150 | $N_y$ | 150 | $N_y$ | 150 |
| $k_{in}$ | 8 | $k_{in}$ | 8 | $W_{max}^{in}$ | 5 |
| $k_{out}$ | 8 | $k_{out}$ | 8 | $W_{max}^{out}$ | 5 |
| $d_0$ | 30 | $d_0$ | 8 | $d_0$ | 20 |
| $n_{draw}$ | 5 | $\eta$ | 0.00025 | $w_{max}$ | 4 |
| | | $r_0$ | 0.2 | $\epsilon$ | $2.15 \times 10^{-4}$ |
| | | T | 10 | $\eta$ | 5 |

### Greedy wiring process principle

The greedy wiring process principle is a normative algorithm that results in formed networks that have the same features as the equilibrium formed networks obtained by running any of the three growth rules described above. This principle provides a direct algorithmic description of which neurons should be connected, without the need for synaptic activity propagation and explicit spontaneous activity.

We begin as for the growth rules above: with a semi-elliptical LGN placed above a semi-infinite unparcellated cortical sheet, and with LGN neurons whose in-degree has already been saturated by inputs from the retina.

We define the initial ‘current’ area to be LGN. The greedy wiring process principle dictates that the shortest possible connections are placed first. We computationally iterate over increasing connection lengths *nΔd* starting from *n* = 0. We begin in the current area, and place all connections from neurons *i* to *j* that satisfy all of the following: (a) neuron *i* is in the ‘current’ layer; (b) neuron *i* is not out-degree saturated; (c) neuron *j* is not in-degree saturated; (d) the Euclidean distance from neuron *i* to neuron *j* is between (*n* − 1/2)Δ*d* and (*n* + 1/2)Δ*d*. This continues with increasing *n* until all neurons in the current layer have become out-degree saturated.

The set of neurons that receive a direct input from the now-out-degree saturated current layer become the new ‘current’ layer. The above process repeats again starting with *n* = 0 until the new ‘current’ layer is out-degree saturated, leading to the formation of successive layers.

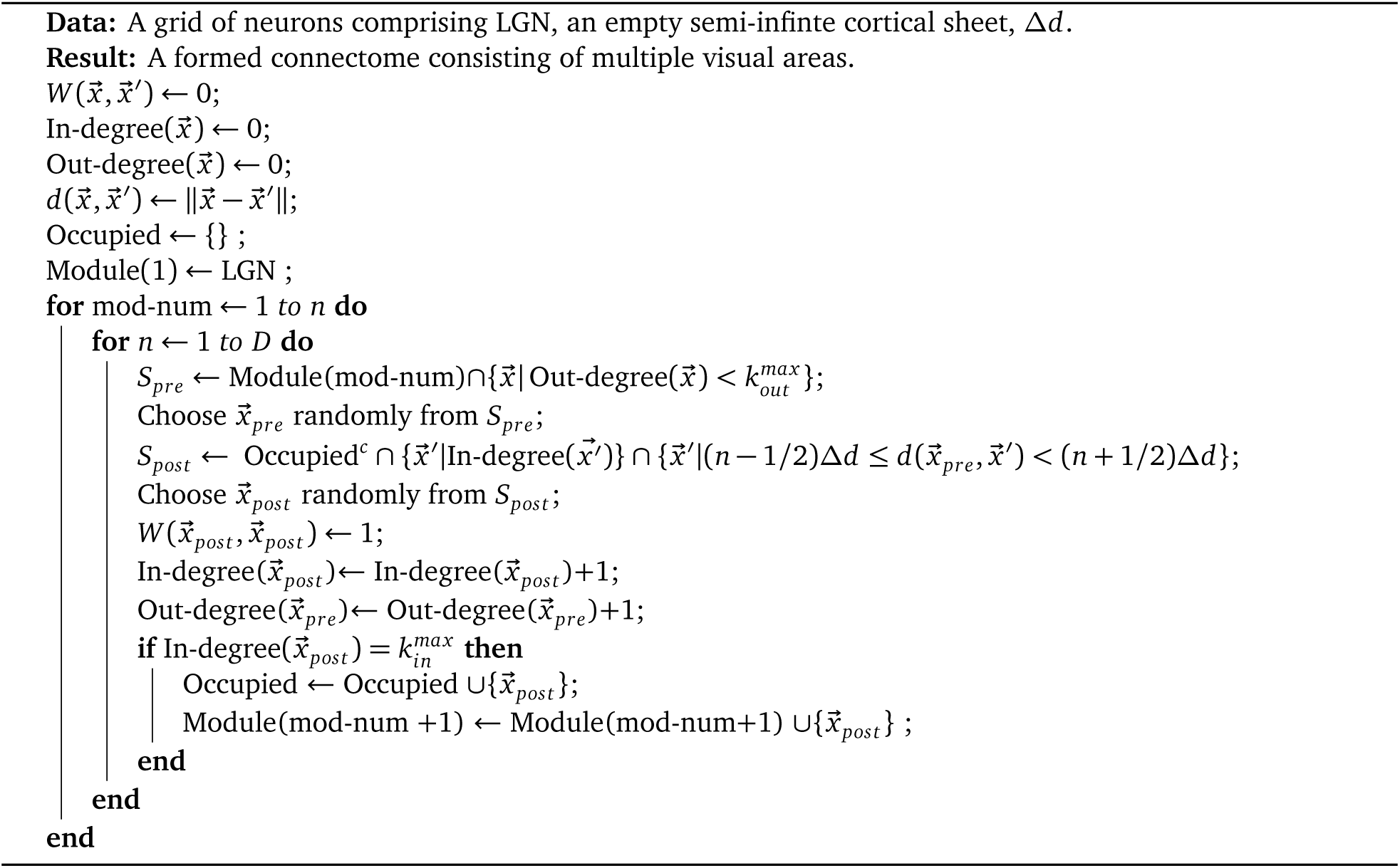
**Algorithm 1**: Greedy wiring process principle

For notational simplicity, the above pseudocode does not make an explicit distinction between *W* and *W_LGN_*. It instead treats all interactions to be encompassed within the same *W* (**x**, **x***^′^*) — the distinction between LGN neurons and those in the cortical sheet can be assumed to lie in the coordinates **x** itself, with LGN neurons being placed at a ε *≪ Δd* distance above the cortical sheet.

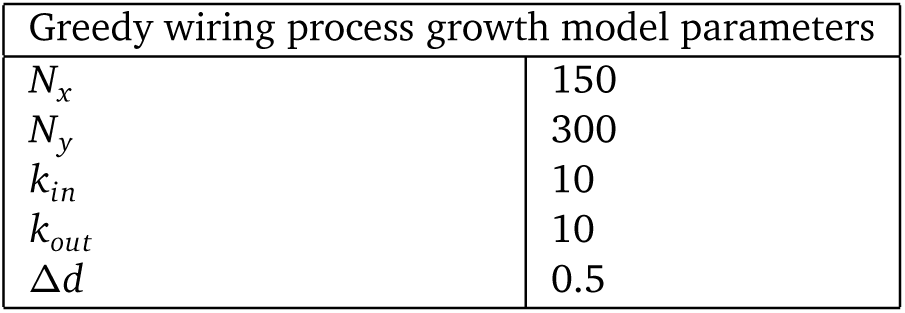

| Greedy wiring process growth model parameters |  |
| --- | --- |
| $N_x$ | 150 |
| $N_y$ | 300 |
| $k_{in}$ | 10 |
| $k_{out}$ | 10 |
| $\Delta d$ | 0.5 |

### Two cell types that generate feed-forward and local connections

In SI Fig. S5, we observed that formed connections in the heterosynaptic competition rule (i.e., whether they are feed-forward connections or local lateral connections) are determined via either of the two parameters, *d*_0_ or γ. We observed that it could be possible to model two co-existing cell types by allowing them to have different values of *d*_0_ or γ. To explore the formation of the visual hierarchy with both cell types, we adapt the greedy wiring minimization process to accommadate both types of cells.

We add a type 2 neuron to the greedy wiring minimization process as follows: Allowing type 2 neurons to throw out local lateral connections would require the neuron to get selected as a presynaptic neuron *before* all other local laterally situated neurons in the same level of ‘hierarchy’ have become in-degree-saturated. We model this by allowing newly connected neurons, if they are of type 2, to immediately get selected to throw connections to the closest available in-degree-unsaturated neurons during the locally greedy wiring algorithm. More specifically, we ignore the hierarchy constraint in selecting neurons in the inner-most for loop, i.e., *S_pre_* is now defined as 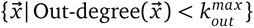.

Note that this is qualitatively similar to what is achieved by increasing γ in the proposed biologically plausible growth rules — a larger γ leads to neurons later in the hierarchy being more active, effectively reducing the distinguishing effect of hierarchy in the network as it forms. Thus, neurons in later areas are more equivalent to earlier neurons for larger γ. Hence, in the locally greedy wiring minimization algorithm, allowing later neurons to be selected ignoring hierarchy qualitatively captures the nature of type 2 neurons as induced by variation of γ (or variation of *d*_0_, cf. Fig. S5).

Thus, this provides a direct, computationally faster method within the greedy wiring minimization principle to recreate the effect of the two discovered cell-types in the biologically plausible learning rules: Cell type 1, follows the usual greedy wiring process principle, and Cell type 2 follows the changed principle described above, allowing Cell type 2 neurons to be included in *S_pre_* as soon as they are connected to the growing network of neurons. We then generate the results of Fig. 9 by creating a cortical sheet with an interleaved mixture of both cell types, and then allow the greedy wiring process algorithm to run until the formation of multiple layers.

## Supplementary Information

**Figure S1:**
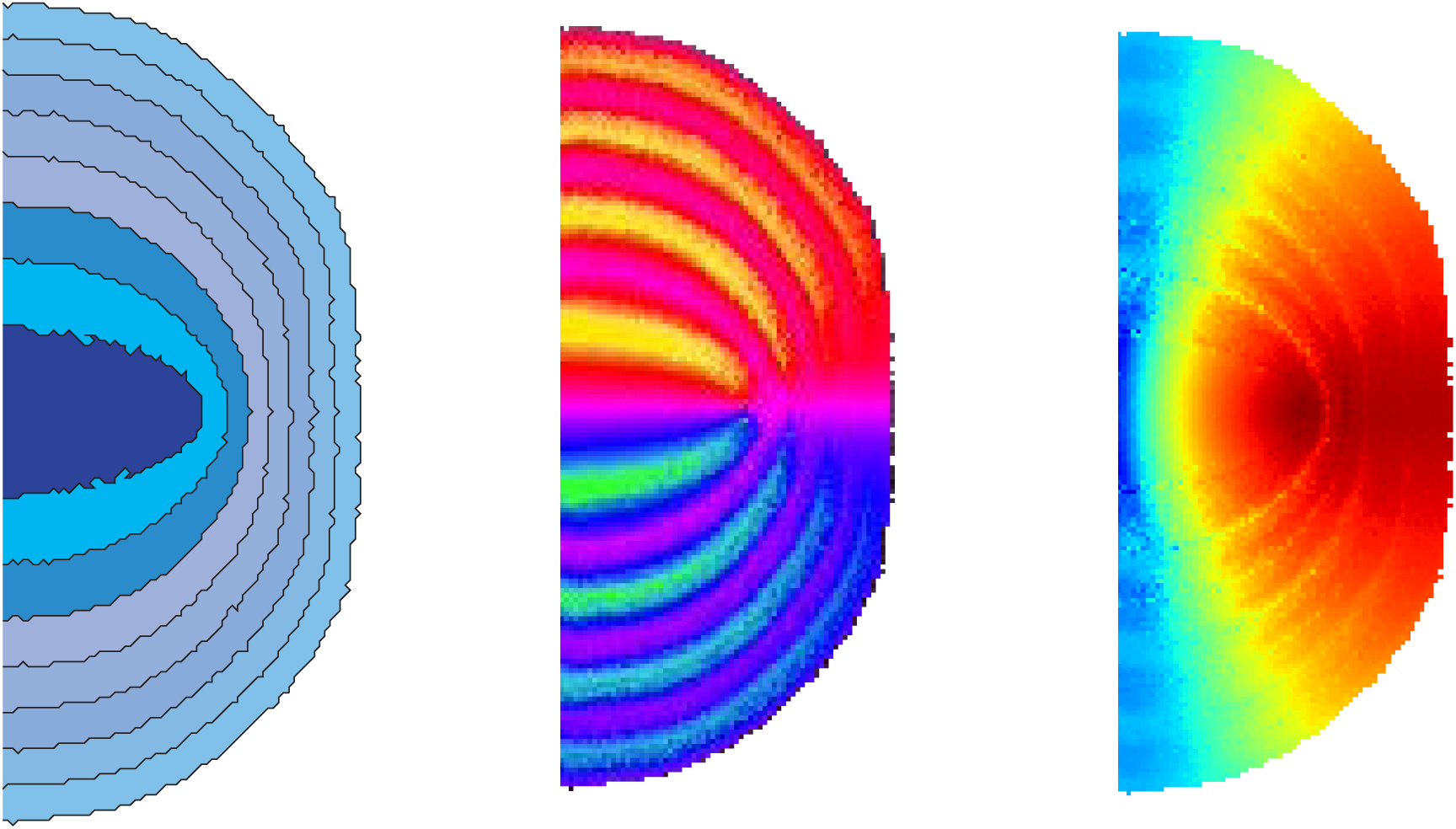
Growth proceeds until the available cortical sheet is exhausted: If the network growth rules are allowed to continue, then the growth continues to form additional hierarchical areas. Each formed area is concentrically arranged around the previous area (left), and all areas exhibit retinotopic mirror-reversed maps (middle: polar angle, right: eccentricity).

**Figure S2:**
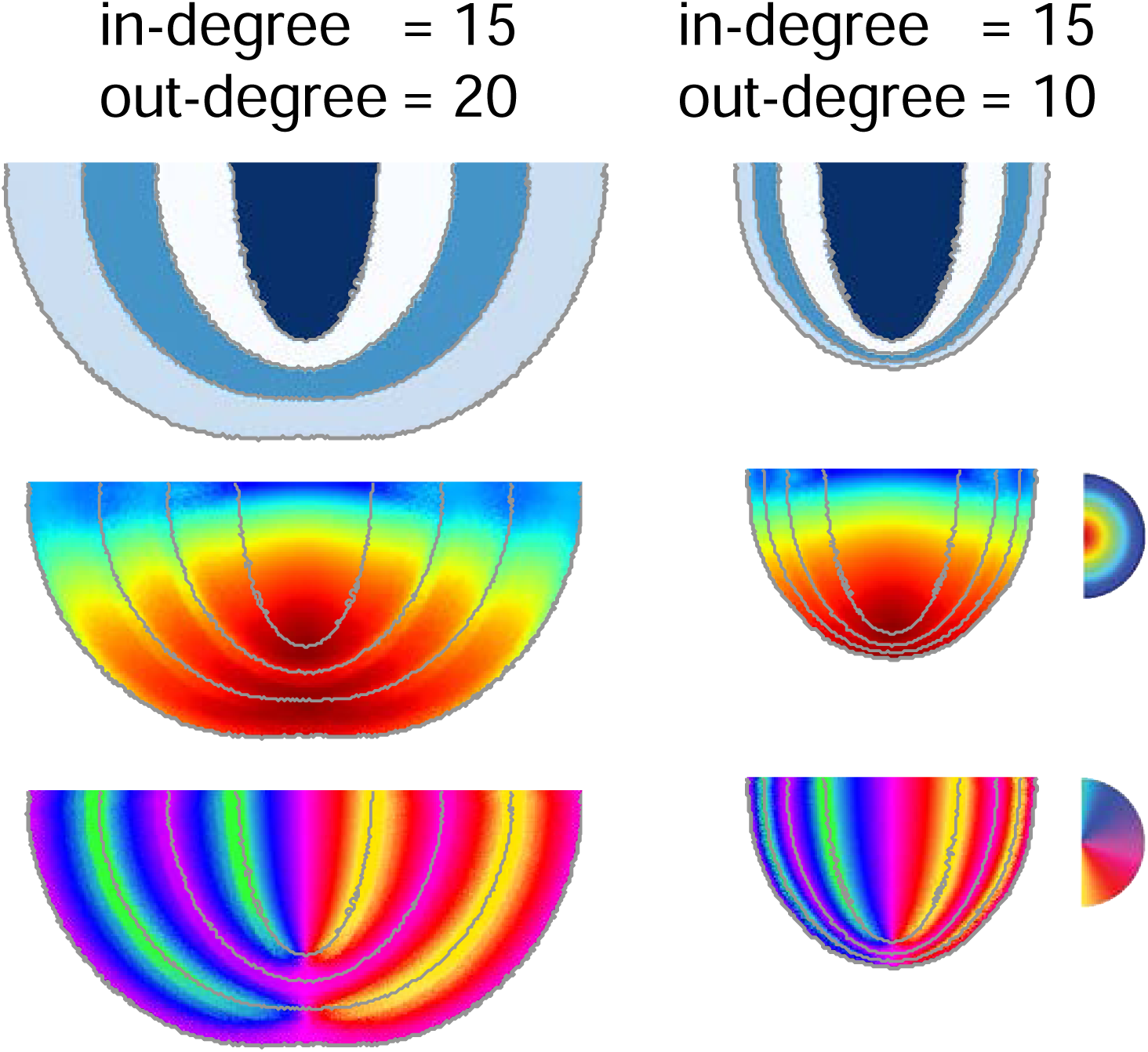
Relative sizes of areas are determined by the in-degree and out-degree of neurons. The ratio of sizes of successive areas is given by the ratio of the out-degree to the in-degree of neurons in the cortical sheet. A ratio larger than 1 (left) results in successive areas increasing in size, and a ratio smaller than 1 (right) results in areas decreasing in size.

**Figure S3:**
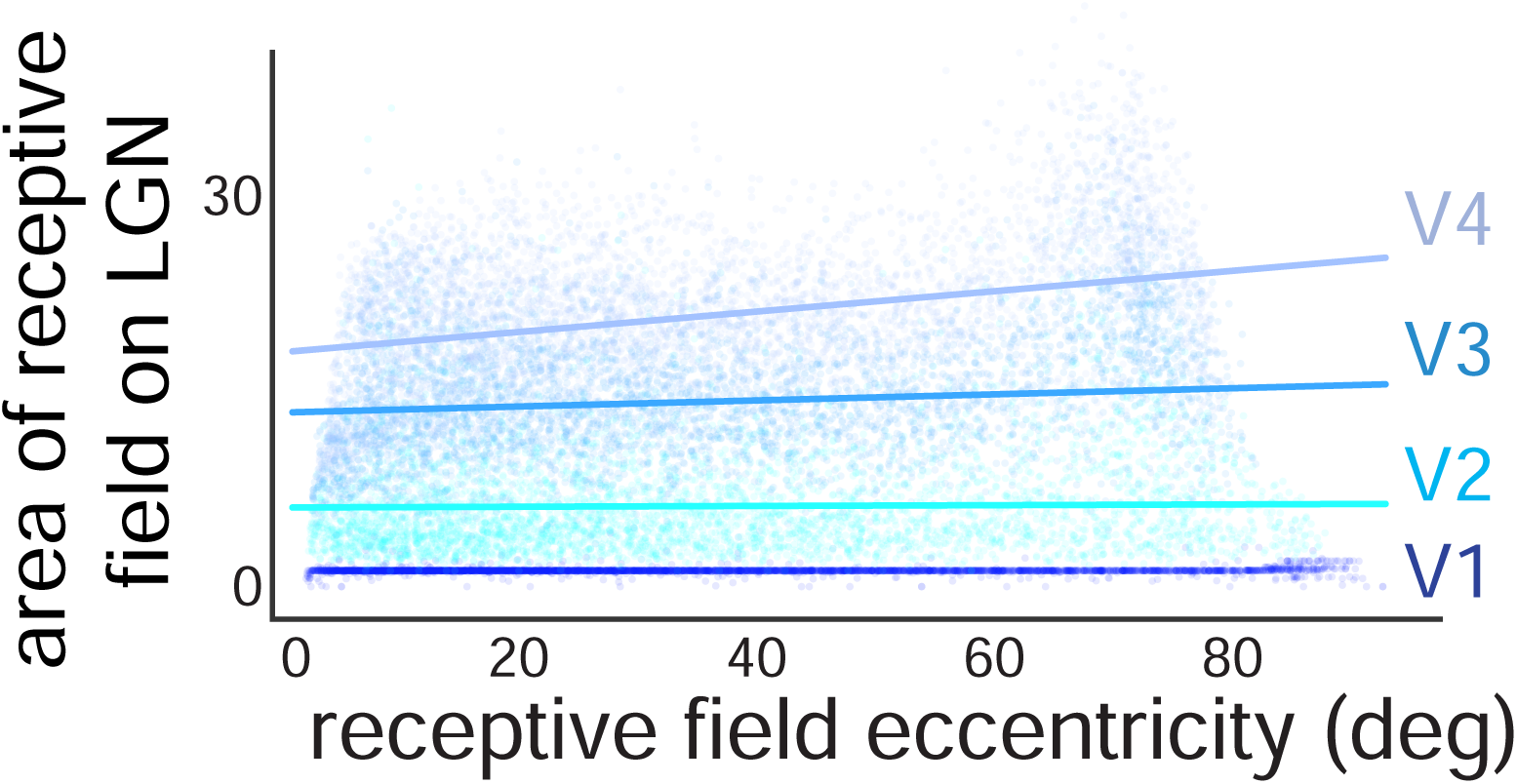
Receptive fields on LGN in neural space do not have eccentricity dependence. Area of receptive field onto LGN for each neuron in the cortical sheet is computed through fitting a two-dimensional Gaussian and computing the product of the two standard deviations. This area shows no significant dependence on the eccentricity of the receptive field in visual field coordinates.

**Figure S4:**
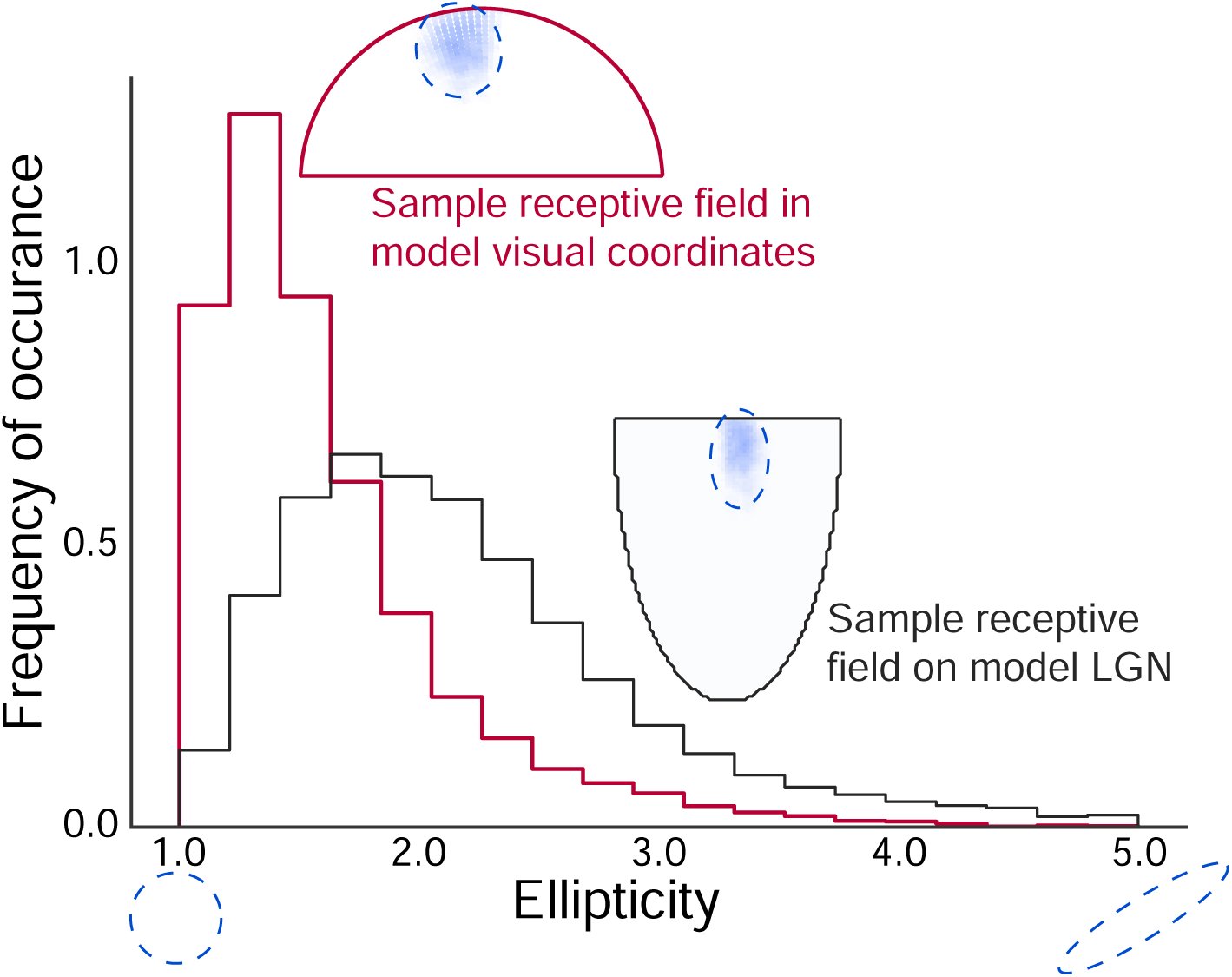
Receptive fields in visual space are more circular than fields in neural coordinates on LGN. A two-dimensional Gaussian function is fit to each receptive field (in visual or neural coordinates). The ellipticity is defined as the ratio of the larger standard deviation to the smaller standard deviation of this Gaussian. The distribution of ellipticities in visual coordinates (in red) tends towards smaller values (and hence more circular fields) as compared to the distribution of ellipticities in neural coordinates (in black). Insets show an example of a receptive field in visual and neural coordinates for a randomly chosen neuron

**Figure S5:**
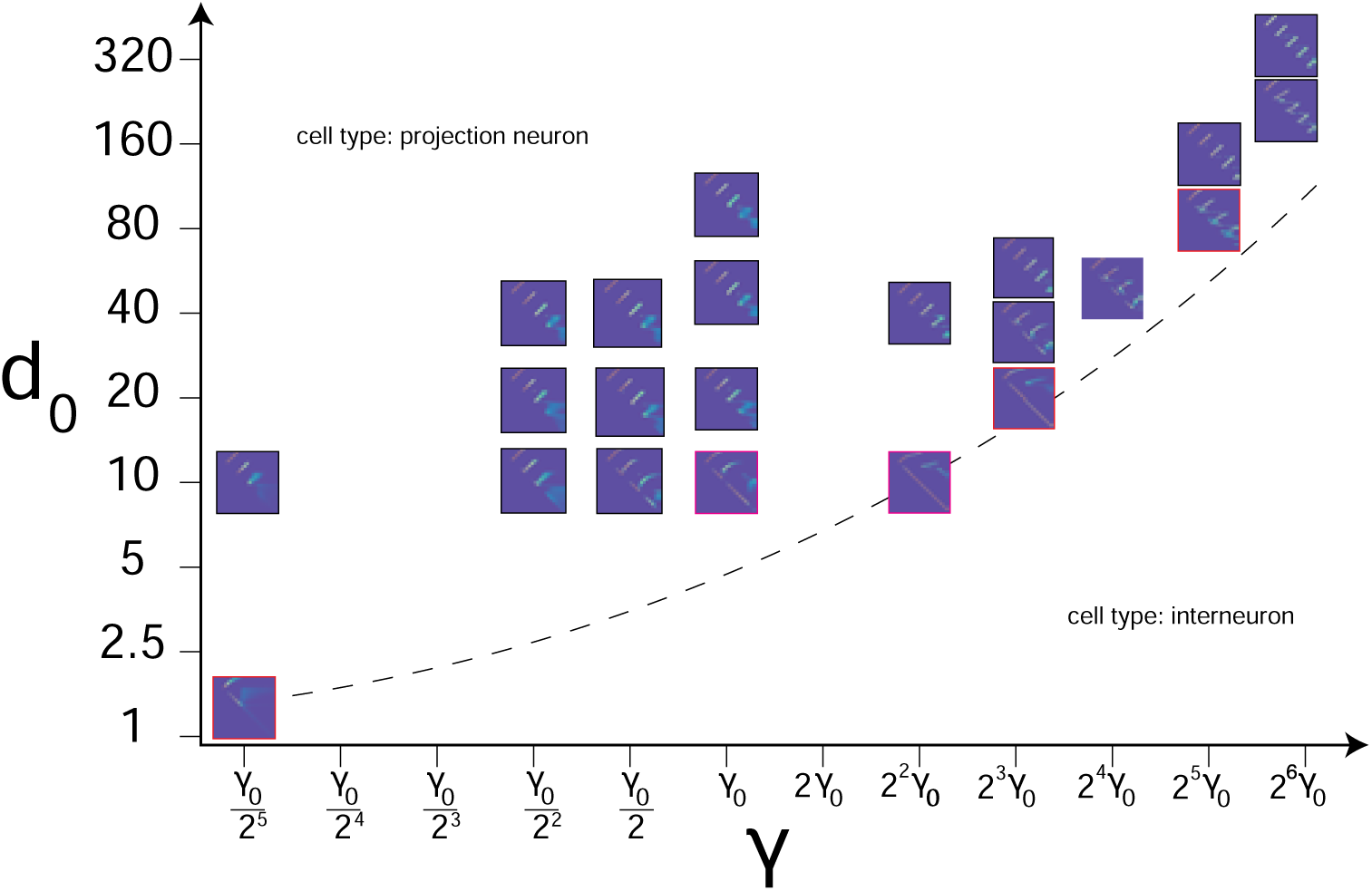
Robustness of the heterosynaptic competition rule to parameters *d*_0_ and γ: Shown is *d*_0_ varied across 2 orders of magnitude and γ across 4 orders of magnitude. Each plot is a 1d simulation and the formed weight matrix is displayed. The simulations shown in red-boxes have formed locally recurrent-like connections, as can be seen by the large number of nonzero elements along the diagonal of the weight matrices. Both axes are on a log scale.

**Figure S6:**
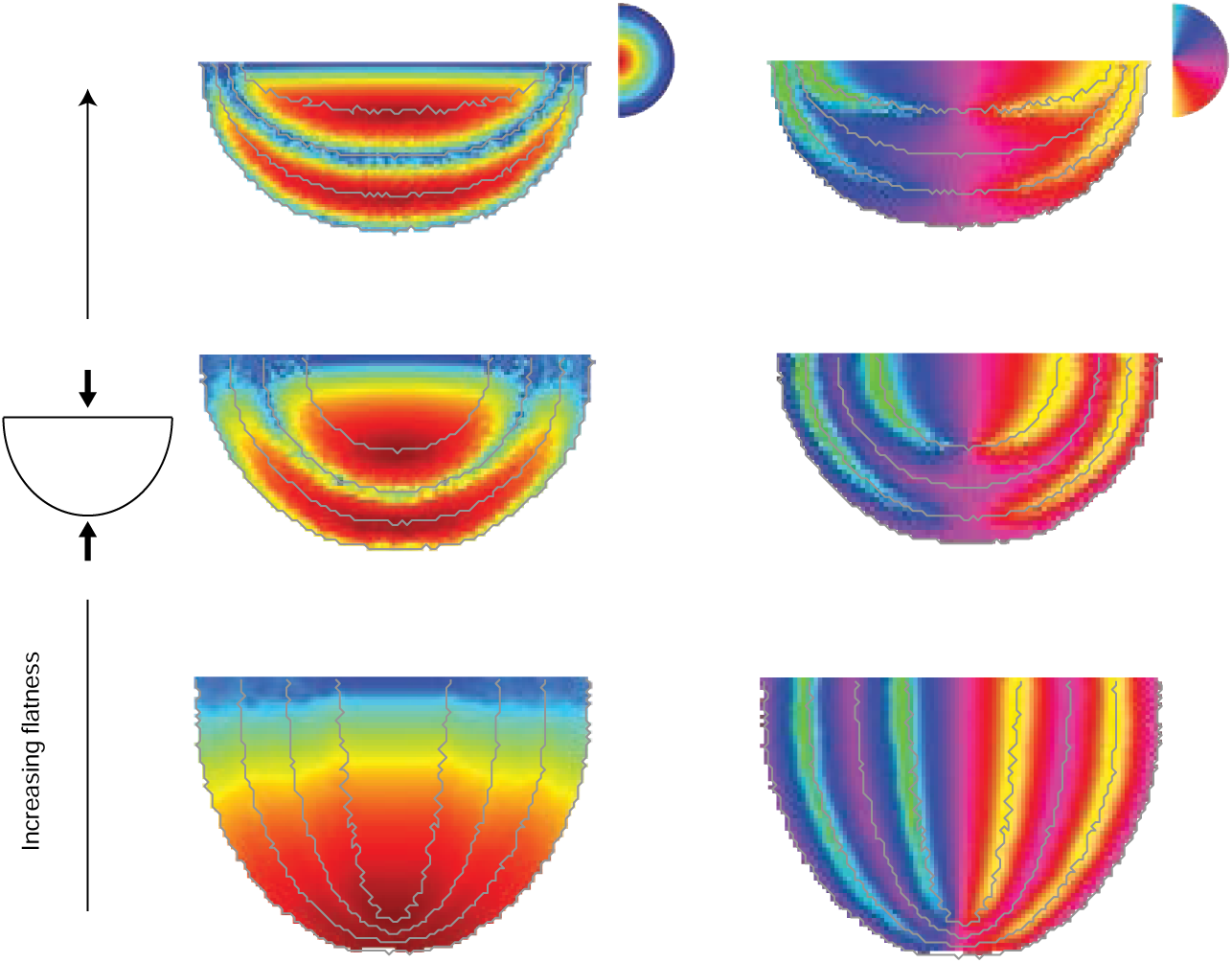
The effect of geometry of LGN on the form of the retinotopic map: (Bottom to top) Increasing flatness of the projection from retina to the putative LGN can lead to retinotopic maps that have alternations in eccentricity as well. Grey lines denote boundaries of areas ascertained by connectivity (which coincides precisely with alternations).

## Notes

### Competing Interest Statement

The authors have declared no competing interest.

